# Inter-subunit Crosstalk via PDZ Synergistically Governs Allosteric Activation of Proapoptotic HtrA2

**DOI:** 10.1101/2021.10.04.462974

**Authors:** Aasna L. Parui, Vandana Mishra, Shubhankar Dutta, Prasenjit Bhaumik, Kakoli Bose

## Abstract

Mitochondrial serine protease – High temperature requirement A2 (HtrA2), is associated with various diseases including neurodegenerative disorders and cancer. Despite availability of structural details, the reports on HtrA2’s mechanistic regulation that varies with the type of activation signals still remain non-concordant. To expound the role of regulatory PDZ domains in promoting synergistic coordination between HtrA2 subunits, we generated heterotrimeric HtrA2 variants comprising different numbers of PDZs and/or active-site mutations. Sequential deletion of PDZs from the trimeric ensemble significantly affected its residual activity in a way that proffered a hypothesis advocating *intermolecular* allosteric crosstalk via PDZ domains in trimeric HtrA2. Furthermore, structural and computational snapshots affirmed the role of PDZs in secondary structural element formation and coordinated reorganization of the N-terminal region and regulatory loops. Therefore, apart from providing cues for devising structure-guided therapeutic strategies, this study establishes a physiologically relevant working model of complex allosteric regulation through a multifaceted *trans*-mediated cooperatively-shared energy landscape.

## INTRODUCTION

Cells respond to different environmental cues by channelizing distinct signal transduction pathways, often controlled by conformational adaptations of multidomain allosteric enzymes. The modulations include subtle loop movements and/or larger domain reorientations upon ligand binding at the allosteric pocket, leading to enhanced substrate catalysis [1, 2]. Interestingly, these conserved regulatory domains that confer a distinct spatial scaffold to the overall structure have been evolutionarily tuned to perform specific functions in several unrelated proteins by accommodating themselves to global structural plasticity in diverse macromolecular milieu [3, 4].

PDZ (Postsynaptic density-95/Discs large/Zonula occludens-1) domains are modular protein-protein interaction moieties specialized for binding to C-terminal motifs of various proteins [5–7]. The multiple ligand-binding sites in PDZ contribute toward their unparallel conformational plasticity relay signals from a distal binding region to the functional site. Human HtrA2 (High temperature requirement protease A2), a PDZ-bearing mitochondrial proapoptotic serine protease that is associated with neurodegeneration and cancer, is an important therapeutic target [8, 9]. Despite exhibiting evolutionarily conserved higher-order architecture with one or two PDZs and a well-defined active site pocket [10], prevalence of low sequence identity and ligand-induced distinct structural signatures provide the basis for the specificity and functional diversity in the HtrA family [9, 11]. One of the unique features of HtrA2 is removal of N-terminal transmembrane region upon apoptotic induction. This leads to its cytoplasmic translocation and subsequent cleavage of inhibitor of apoptosis proteins (IAPs) through an exposed AVPS motif thus promoting cell death through a binary activation mode [12]. Furthermore, HtrA2 promotes non-classical caspase-independent apoptosis [13, 14] via its intricate serine protease activity. However, the complex mechanism through which it promotes initiation of apoptotic cascades or regulates protein quality control remains elusive.

The crystal structure of the catalytically inactive apo-HtrA2 [15] proposed a mechanistic model wherein the otherwise catalytically competent protease domain are stereochemically shielded by PDZ from the same monomer in its basal state thus implicating a *cis*-mediated inhibitory role of PDZs. However, subsequent reports that demonstrated a significant decrease in the protease activity of a PDZ-lacking variant [16, 17] as well as multiple activation routes of HtrA2 [18–21] challenged this pre-existing tenet. Revisitation of the mode of HtrA2 activation highlighted a dual regulatory inter-molecular switch (through allosteric ligand-binding to PDZs or N-terminus) promoted coordinated structural reorganizations at the regulatory loops essential for HtrA2 activation [17, 22]. However, these studies as well as multiple structural reports [15, 18, 23] fail to elucidate the intricate real-time conformational dynamics with respect to different modes of activation viz. substrate binding or temperature thus leaving a huge vacuity toward understanding the multifaceted role of HtrA2.

Evolutionarily, PDZs are known to govern the active and/or inactive state(s) of HtrA oligomers. Apart from the N-terminal-mediated regulation in HtrA2 [22], the transition from apo-form to the proteolytically active state seems to be strictly governed by PDZs, owing to its specific interactions with numerous C-terminal binding partners such as cytoskeletal proteins and other apoptosis-related molecules. The basic requirement of HtrA proteases to exist as large oligomeric ensembles, despite having all the functional elements present in a single subunit, is quite astonishingly profound. The aim of this study is to recognize the contribution of each HtrA2 subunit towards the dynamic energy landscape within the trimeric ensemble and to identify how individual PDZs, besides allosterically regulating HtrA2 function, help attain an active functional state upon multimodal stimulation. To achieve this, we generated different heterotrimeric HtrA2 variants that lack one or two PDZ domains and performed comprehensive enzyme kinetic analyses of each variant with a C-terminal binding partner. Through active-site chemical profiling [24], we established that binding of substrate to a single PDZ not only significantly affects the proteolytic activity of adjacent subunits, but also compensates for the loss of active-sites within the ensemble. In case of both substrate- and temperature-based activations, we demonstrated an intricate predominant interdomain networking (*trans* PDZ-protease crosstalk) that leads to subtle reorganization of the active-site loops and formation of a protease-substrate complex that is tuned for efficient catalysis. This networking is responsible for maintaining the overall proteolytic activities of each HtrA2 subunit via cooperatively-coupled energy dynamism within the trimeric ensemble. Furthermore, this study emphasizes the importance of the evolutionarily conserved PDZ in determining the active functional state of HtrA2 that is catalytically more competent under a specific cellular scenario and is also energetically most favorable. The concept encompassing such functionally important events, which can be transmitted long-range to affect the affinity or catalytic efficiency of a distant active-site, poses a challenge and necessity to identify the communication pathway underlying the functional effects of such larger macromolecular assemblies within the cell.

## RESULTS

### Generation of heterotrimeric HtrA2 variants with different number of PDZ domains

For understanding the specific function of PDZs in propagating allosteric communication, we generated heterotrimeric variants of HtrA2, which differed in the number of PDZs. The constructs used include the homotrimeric His_6_-tagged wildtype (WT or WWW) and the His_3_-tagged ΔPDZ construct (N-SPD or ΔΔΔ) (Fig. 1A). The purified homotrimers were mixed in equimolar concentrations and subjected to mild denaturing conditions. Subsequent renaturation led to the formation of four plausible HtrA2 variant combinations - two homotrimers (WWW, ΔΔΔ) and two heterotrimers (WWΔ, WΔΔ) (Fig. 1A). The additional N-terminal FLAG-tag in ΔΔΔ imparted ∼0.2 difference in pI (as estimated by ProtParam software) [25], thus enabling the variants in the renatured mixture to be well resolved in native PAGE (Fig. 1B). Furthermore, when probed with anti-FLAG antibody, three out of four bands (having Δ subunit) were spotted with varying intensities (different number of FLAG epitopes), confirming the distinctiveness of the heterotrimers (Fig. 1C). These renatured variants were further separated using a modified version of IMAC (Immobilized metal affinity chromatography), based on the differential affinity toward histidine tag. As W and Δ subunit differed in the number of C-terminal histidine tag residues, batch purification with a narrow imidazole gradient led to the elution of ΔΔΔ in flow-through, followed by WΔΔ, WWΔ, and WWW eluting between 50-250 mM imidazole gradient (Fig. 1D). The heterotrimers obtained were stable, as estimated by native PAGE (Fig. 1E). The purity of WWΔ and WΔΔ were found to be slightly lower (∼90%) as compared to the other variants due to interference from the adjoining variant. For further enzymatic studies, the homotrimers were purified, denatured and renatured separately.

**Figure 1:**
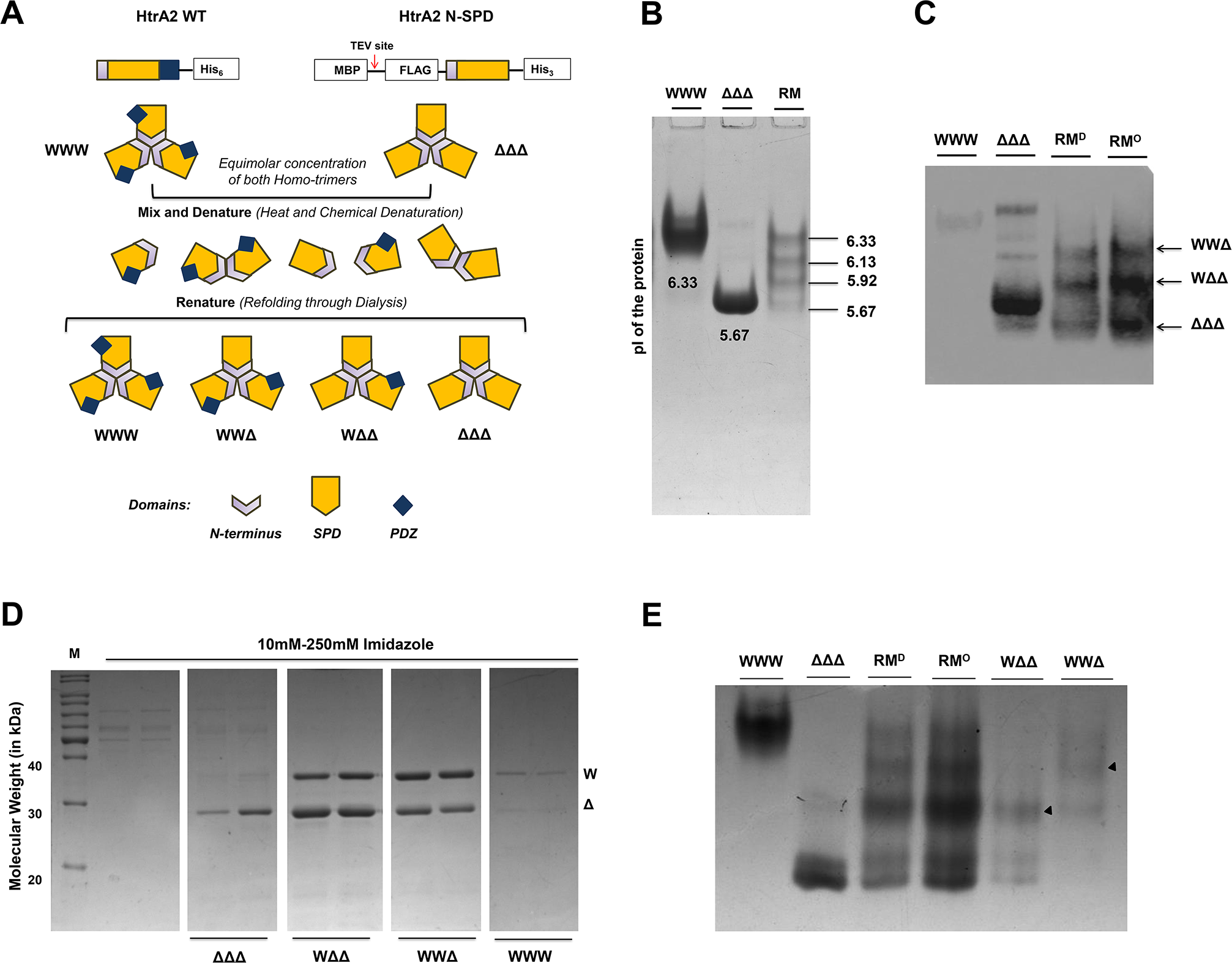
HtrA2 variants with one or two PDZ domains. **(A)** Schematic model depicting the denaturation-renaturation strategy that is used to generate heterotrimeric HtrA2 variants (WWΔ and WΔΔ). **(B)** Native PAGE showing generation of precisely four combinations of HtrA2 variants. Due to significant differences in pI, the electrophoretic separation of the four HtrA2 variant combinations in the renatured reaction mixture (RM) was observed in comparison to the parent homotrimers-wildtype (WWW) and PDZ-deleted construct (ΔΔΔ). The concentration of the homotrimers used in this analysis was same as that used in the denaturation process to check the presence of any higher order oligomer formation at this concentration. **(C)** Western Blot analysis confirming the identity of heterotrimers. Native PAGE, as described in (B), was used for transfer of the proteins and subsequent blotting analysis. As the parent ΔΔΔ construct consisted of an N-terminal FLAG tag, the trimers containing Δ subunit (WΔΔ, WWΔ and ΔΔΔ) were easily probed in the reaction mixture using Anti-FLAG antibody. No band was probed in the WWW lane, while presence of few non-specific bands was observed in ΔΔΔ. The renatured reaction mixture was loaded in diluted form (RM^D^) and in the original concentration (RM^O^) to confirm the identity of heterotrimers. **(D)** SDS-PAGE showing the separation of heterotrimeric HtrA2 variants. The variants were separated from the reaction mixture using a very narrow gradient of imidazole in the elution buffer. The separation occurred on the basis of number of histidine residues present in the C-terminal His tag. WΔΔ eluted in the initial gradients, followed by WWΔ and WWW. M: Protein marker. **(E)** Native PAGE showing the separation of heterotrimeric HtrA2 variants. The separated heterotrimeric variants (WΔΔ and WWΔ indicated with filled black triangles in their respective lanes) were identified on the basis of their electrophoretic mobility with respect to their relative position in the reaction mixture (RM^D^ and RM^O^).

### Presence of single or multiple PDZ domains influences HtrA2 proteolytic activity

To understand the effect of PDZ deletion on the overall residual activity and specificity of HtrA2 variants, time-based proteolytic assays were performed using the generic substrate β-casein. This substrate has a putative binding site (GPFPIIV) that interacts with ‘YIGV’ groove of HtrA2 PDZ [16] and mimics the canonical PDZ-mediated allosteric modulation. It was observed that although the HtrA2 variants exhibited similar cleavage pattern (with three distinct bands), the cleavage time differed significantly with removal of each PDZ from the trimer (Fig. 2A). While variants having at least one PDZ (WWW, WWΔ and WΔΔ) cleaved within 10 min of incubation with a serial decline in the cleavage rate with PDZ deletion, ΔΔΔ displayed significant reduction of activity with 100% substrate cleavage only by ∼60 min (Fig. 2B). For quantitative evaluation, fluorometric analysis was performed using varying concentrations of FITC-labeled β-casein as a substrate. Interestingly, a decreasing trend for V_max_ was also observed with removal of each PDZ domain. WWW displayed the highest V_max_, followed by WWΔ and WΔΔ, which demonstrated approximately two- and four-fold decrease respectively, while ΔΔΔ showed a ∼10-fold decrease in V_max_ (Fig. 2C and Table 1).

**Figure 2:**
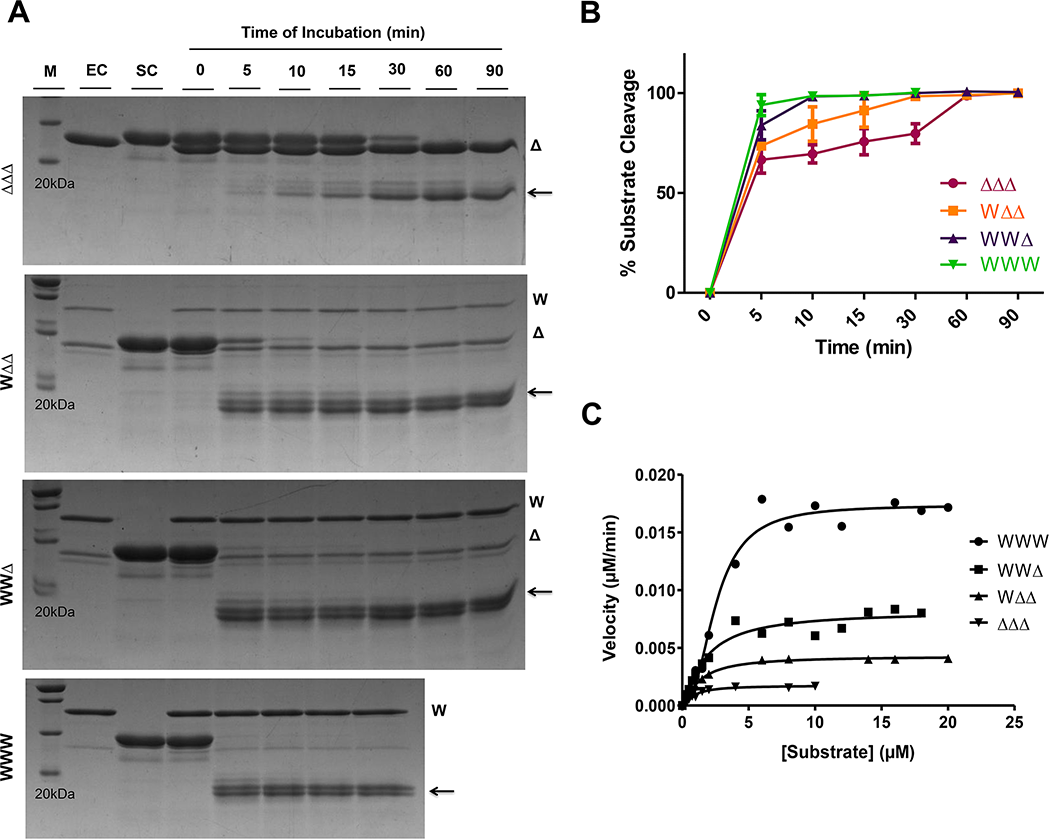
Time-course proteolytic cleavage assays of HtrA2 variants. **(A)** Qualitative gel-based proteolytic cleavage of β-casein by HtrA2 variants. 2 μg of each enzyme variant was incubated individually with 6 μg of substrate β-casein at 37 °C for different time-points between 0-90 min. The reaction at each time point was stopped with Laemmli buffer at 100 °C. Reaction samples were resolved by 12% SDS-PAGE and the cleavage pattern was visualized with Coomassie brilliant blue staining. M: Protein marker; EC: Enzyme control; SC: Substrate control. Arrows indicate the cleaved products. **(B)** Semi quantitative analysis of HtrA2 protease assays. Percentage substrate cleaved at the end of each reaction time point was calculated by quantifying the intensity of residual β-casein at each time point relative to the uncleaved substrate control (SC) using *GelQuant.NET* software and was plotted using Graphpad Prism software. WWW displayed maximum substrate cleavage activity within 5 min, followed by WWΔ and WΔΔ that cleaved 90% substrate by 5 and 10 min respectively. ΔΔΔ displayed 100% substrate cleavage between 30-60 min. The error bars represent the standard errors (SE) calculated from three independent experiments. **(C)** Quantitative assessment of the HtrA2 proteolytic activity. Initial velocities V_0_ of HtrA2 and its variants were calculated using FITC labeled–β casein as the substrate. The solid lines are the nonlinear least squares fit of the data to the Hill form of the Michaelis–Menten equation: Velocity = V_max_/[1 + (K_0.5_/[substrate])^n^] using Graphpad Prism software.

**Table 1:** Steady state enzyme kinetic parameters for HtrA2 variants. The initial rates for the cleavage of FITC labeled–β casein by different HtrA2 variants were measured and fitted to the Hill form of the Michaelis–Menten equation to determine the steady-state kinetic parameters (Figs 2 and S2). Values are the mean ± SEM and are generated from data points obtained from at least three independent experiments.

| **HtrA2**<br>**Variants** | **$V_{\max}$ ($\text{Ms}^{-1}$)**<br>**$\times 10^{-10}$** | **$K_{0.5}$ ($\mu\text{M}$)** | **$k_{\text{cat}}$ ($\text{s}^{-1}$)**<br>**$\times 10^{-05}$** | **$k_{\text{cat}}/K_m$ ($\text{M}^{-1} \text{s}^{-1}$)** | **Hill constant** |
| --- | --- | --- | --- | --- | --- |
| **WWW** | $2.892 \pm 0.0002$ | $2.513 \pm 0.2$ | 14.46 | 57.534 | $2.357 \pm 0.3$ |
| **WWΔ** | $1.360 \pm 0.0007$ | $1.738 \pm 0.4$ | 6.81 | 39.154 | $1.231 \pm 0.3$ |
| **W$\Delta\Delta$** | $0.713 \pm 0.0001$ | $1.293 \pm 0.1$ | 3.57 | 27.578 | $1.272 \pm 0.1$ |
| **$\Delta\Delta\Delta$** | $0.289 \pm 0.0001$ | $0.869 \pm 0.1$ | 1.44 | 16.607 | $1.367 \pm 0.3$ |
| **W<sub>I</sub>$\Delta\Delta$** | $0.569 \pm 0.00008$ | $0.931 \pm 0.1$ | 2.84 | 30.546 | $1.824 \pm 0.2$ |
| **W<sub>I</sub>W<sub>I</sub>Δ** | $1.219 \pm 0.0003$ | $1.771 \pm 0.2$ | 6.09 | 34.401 | $2.353 \pm 0.5$ |
| **WΔ<sub>I</sub>Δ<sub>I</sub>** | $0.157 \pm 0.00004$ | $0.423 \pm 0.1$ | 0.78 | 18.504 | $2.714 \pm 0.9$ |
| **WWΔ<sub>I</sub>** | $0.412 \pm 0.0002$ | $0.877 \pm 0.2$ | 2.11 | 24.022 | $2.077 \pm 0.8$ |

Although PDZ deletion resulted in only minor changes in the positive co-operativity of heterotrimers, the apparent sequential decline in V_max_ proposes a strong possibility of inter-subunit crosstalk between PDZ and adjacent protease domain, which eventually influences the second step of enzyme kinetics i.e. substrate catalysis. Alternatively, ΔΔΔ exhibited maximum substrate binding affinity (K_m_), followed by WΔΔ and WWΔ. This is expected as removal of PDZ inhibition is known to open up the base of the pyramidal ensemble [15, 26], thereby increasing substrate accessibility leading to pseudo-affinity exhibited by enzyme. Despite this, we observed a marked decrease in V_max_ and substrate turnover (k_cat_) rates for WWΔ, WΔΔ and ΔΔΔ. These results hint toward the presence of a malformed oxyanion hole in PDZ-lacking variants that also corroborates well with our previous studies [16, 17]. To assess the relative orientation of oxyanion hole residues, comparative 100 ns MDS analyses of these Δ-containing variants were performed, keeping WWW as the reference structure. In-depth structural comparison revealed a 90° anti-clockwise flip in the imidazole ring of F303 away from the catalytic triad residue H198 for WWΔ, WΔΔ and ΔΔΔ (Fig. S1). This distal movement disrupts the nucleophilic exchange between F303 and H198 that is indispensable for stabilizing the negatively charged intermediates generated due to H198-induced deprotonation of the catalytic serine S306 [9, 16]. This further explains the decrease in catalytic efficiencies, with ΔΔΔ showing approximately four-fold, while WΔΔ and WWΔ approximately two- and ∼1.5-fold decrease respectively, as compared to WWW. Overall, these results suggest that all the three PDZs of the HtrA2 trimeric ensemble govern its proteolytic activation by maintaining proper orientation of the catalytic triad, which is a prerequisite for a stabilized transition state formation and subsequent substrate catalysis.

### Catalytic pocket reorientation and structural perturbation captured through MDS

To investigate the role of PDZ in modulating substrate accessibility, *in silico* docking analysis was performed using β-casein peptide (GPFPIIV) and HtrA2 variants. Comparison of the apparent binding affinities showed ΔΔΔ having higher docking score than WWΔ and WΔΔ (Table S1). Interaction analysis of the docked complexes revealed the propensity of the peptide to bind with H394, P409 and G410 of PDZ (Figs. S2A, S2B and S2C). However, due to higher accessibility in ΔΔΔ, the peptide directly forms interactions with the active-site residues (H198, D228 and S306), resulting in higher binding score compared to the PDZ-containing variants (Fig. S2D). Furthermore, to discern the underlying mechanism contributing to differences in catalytic efficiency, MDS analyses were performed with these docked complexes. Previous reports showed that HtrA2 undergoes catalytic triad reorientation by moving H198 towards D228, and S306 away from H198, upon activator/substrate binding at the distant PDZ domain [9, 16]. Comparative distance analysis of 1000 ns MDS of the peptide-bound and unbound HtrA2 variants showed similar catalytic triad movement for WWW, resulting in successful opening of the pocket. However, with removal of PDZ, the opening of the catalytic pocket decreased gradually in WWΔ and WΔΔ (Table S1 and S2). Although these movements enabled slight opening of the catalytic pocket for WWΔ and WΔΔ, the opening might not be sufficient to exhibit catalysis as efficiently as WWW and hence supports our aforementioned enzyme kinetic studies. ΔΔΔ, on the other hand, showed closing of the catalytic pocket upon peptide binding, as the distance between H198 and S306 was found to be decreased by 0.4 Å, 0.5 Å and 0.4 Å in chains A, B and C, respectively (Table S2).

Apart from substrate-based activation, the apo-form of HtrA2 is known to be activated by increase in temperature [26], with optimum activation temperature ranging between 45 °C −55°C [17]. To investigate whether HtrA2 undergoes similar catalytic triad reorientation with temperature, the variants were subjected to MDS with temperature elevated to 323 K. Temperature-induced activation also demonstrated a similar trend in opening of the catalytic pocket, with WWW showing the highest deviation of 1.4 Å, whereas for WWΔ and WΔΔ, the maximum deviation was not more than 0.4 Å (Table S2). However, as expected, ΔΔΔ demonstrated reverse movement of the active-site residues upon temperature induction that might result in improper catalytic pocket formation.

To further dissect the underlying conformational changes, metadynamics MDS was performed for the β-casein bound and temperature-afflicted forms of HtrA2 variants that generated free-energy landscape, representing segregated conformations of the systems calculated over a collective set of catalytic triad distances [27]. In both the cases, two separate conformations were evident for WWW, corresponding to their H198-S306 or H198-D228 distances. The two lowest conformations were observed at 1.6 and 1.5 kcal/mol, where the former ΔG value tends to reside at higher catalytic triad residue distances when bound to the activating peptide (Fig. 3A). The two separate lowest conformations of WWW at optimum catalytic pocket distances indicate that opening up is a direct effect of the global conformational changes (Fig. 3B). Similar effect was observed in WWΔ with two discrete conformations at different free energy levels (Figs. 3C and 3D). However, for WΔΔ, though there were a few separately placed conformations, distinct energy barriers among them were missing, which is indicative of transient conformational changes that are not conducive toward enhancing its catalytic efficiency (Figs. 3E and 3F). Furthermore, in ΔΔΔ, single rigid conformation was observed indicating zero transitional state (Figs. 3G and 3H). This structural rigor might be due to the absence of any regulatory domain such as PDZ.

**Figure 3:**
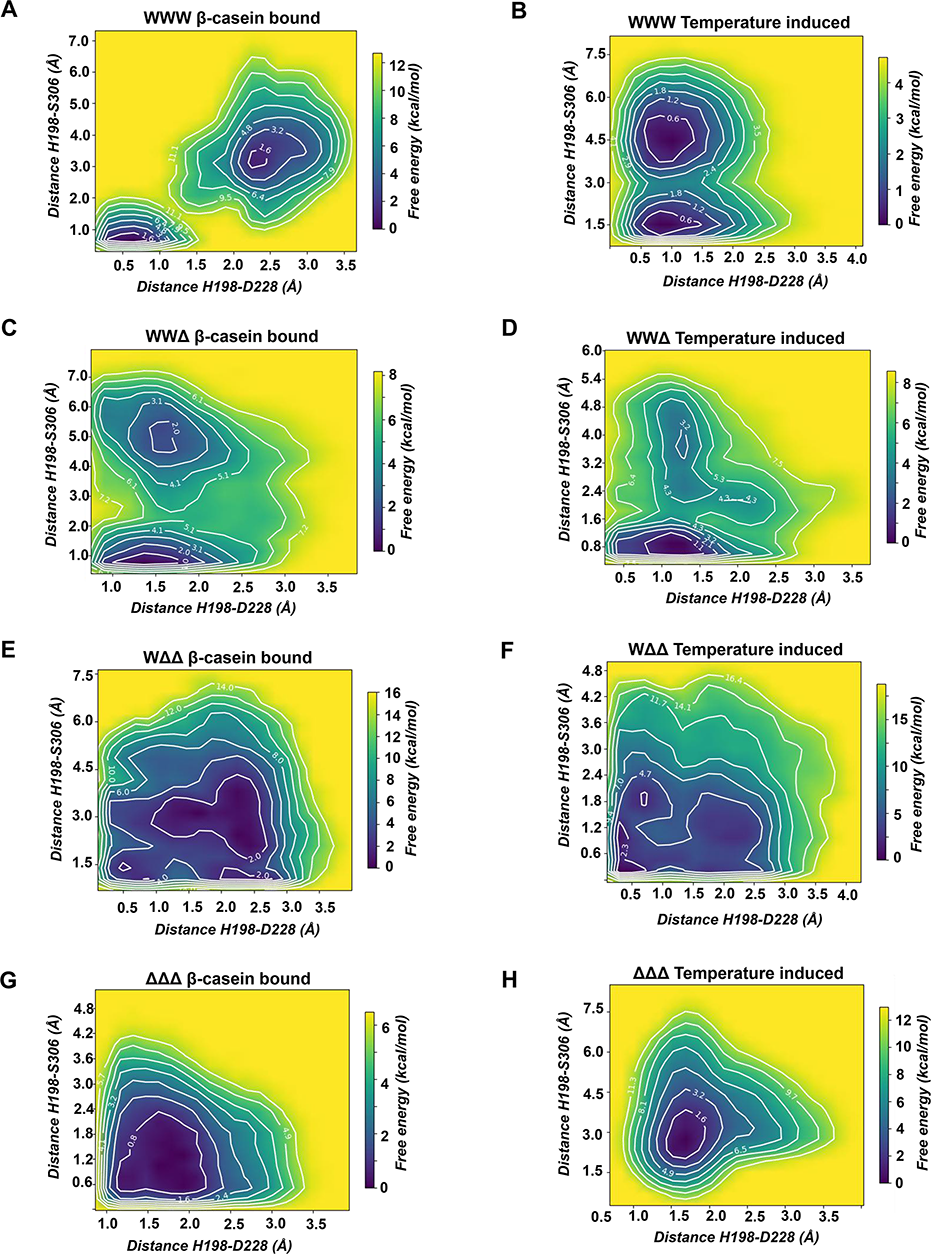
Free energy landscape diagrams of HtrA2 variants generated on the basis of catalytic triad orientation. Contour plots for β-casein bound and temperature-induced **(A, B)** WWW, **(C, D)** WWΔ **(E, F)** WΔΔ and **(G, H)** ΔΔΔ, respectively, where each contour joined all the structures with equal free energies (in kcal/mol). Free energies for all the variants were calculated using the intermolecular distances among the catalytic residues (H198, D228 and S306) as collective variables. The energy gradients have been represented by blue-yellow spectrum where blue represents the lowest and yellow represents the highest free energy.

### Intermolecular PDZ-protease crosstalk governs different HtrA2 activation modes

From the aforementioned studies, it is evident that extensive conformational changes with PDZ movement are imperative for the activation of HtrA2 variants. To understand the role of intra- and/or intermolecular crosstalk in allosteric induction, an activity-based chemical profiling using TAMRA fluorophosphonate (TAMRA-FP) was performed. TAMRA-FP specifically labels the catalytic serine of an active serine hydrolase through covalent modification of the side chain hydroxyl group [24], thus facilitating their detection through the linked fluorescein dye. This study was envisaged to delineate how the two activation modes (substrate- or temperature-induced) influence the active-site reactivity of the attached (*cis*) and/or neighboring (*trans*) protease domain, and the subsequent variations in the conformational dynamics upon activation.

#### Substrate-induced activation

Upon activation, labeling of each subunit (W or Δ) was efficiently determined through fluorescence-based SDS-PAGE. The active-site chemical profiling was performed with PDZ activator-cum-substrate β-casein. As expected, neither WWW nor ΔΔΔ subunits were TAMRA-labeled in their basal states (Fig. S3A). Lack of modification in WWW might be due to over-crowding of PDZs in the apo-form. However, in the unbound WWΔ and WΔΔ variants, W subunit showed ∼2.4- and ∼1.8-fold increase respectively, while Δ subunit exhibited ∼2.8- and ∼5.3 fold respectively as compared to their respective homotrimers; thus suggesting occurrence of favorable conformational changes in and around the active-site of adjacent Δ subunit(s) with removal of PDZ(s) (Fig. S3A and S3C). On the other hand, the bound WWW and ΔΔΔ complexes exhibited highest and lowest fluorescence labeling respectively (Fig. S3A). This confirms that the binding event engages PDZs and greatly alters the activity of the entire trimeric ensemble. Interestingly, Δ subunit of WWΔ showed greater modification than WΔΔ (Fig. S3C). This might be attributed to the simultaneous influence of two PDZs on the active-site of a single Δ subunit through an intricate intermolecular PDZ-protease interplay. In WΔΔ, although the PDZ-crowding is less, the lesser fluorescence equates to modification of two Δ subunits with a single PDZ. These observations ascertain presence of a predominant *trans* PDZ-protease crosstalk between the ligand-bound PDZ of one subunit with the protease domain of the adjacent subunit. For quantitative validation, the TAMRA-FP intensities were compared with respect to their corresponding basal states so as to determine the fold change in active-site modification (Table S3A). While bound WWW represents 11-fold increase in TAMRA fluorescence compared to the unbound form, the W subunit of bound WWΔ and WΔΔ exhibit five- and four-fold increase respectively (Fig. 4A). Interestingly, each Δ subunit of these heterotrimeric variants, WWΔ and WΔΔ, demonstrates increase in labeling by seven- and three-fold respectively compared to their unbound forms (Fig. 4A), thus correlating TAMRA-modification of the Δ subunit on the total number of engaged PDZs. The observed 2.5-fold fluorescence increase in the substrate-bound ΔΔΔ might be attributed to greater accessibility to the active site in absence of PDZs that is in concurrence with the lower K_m_ value earlier obtained for this variant.

**Figure 4:**
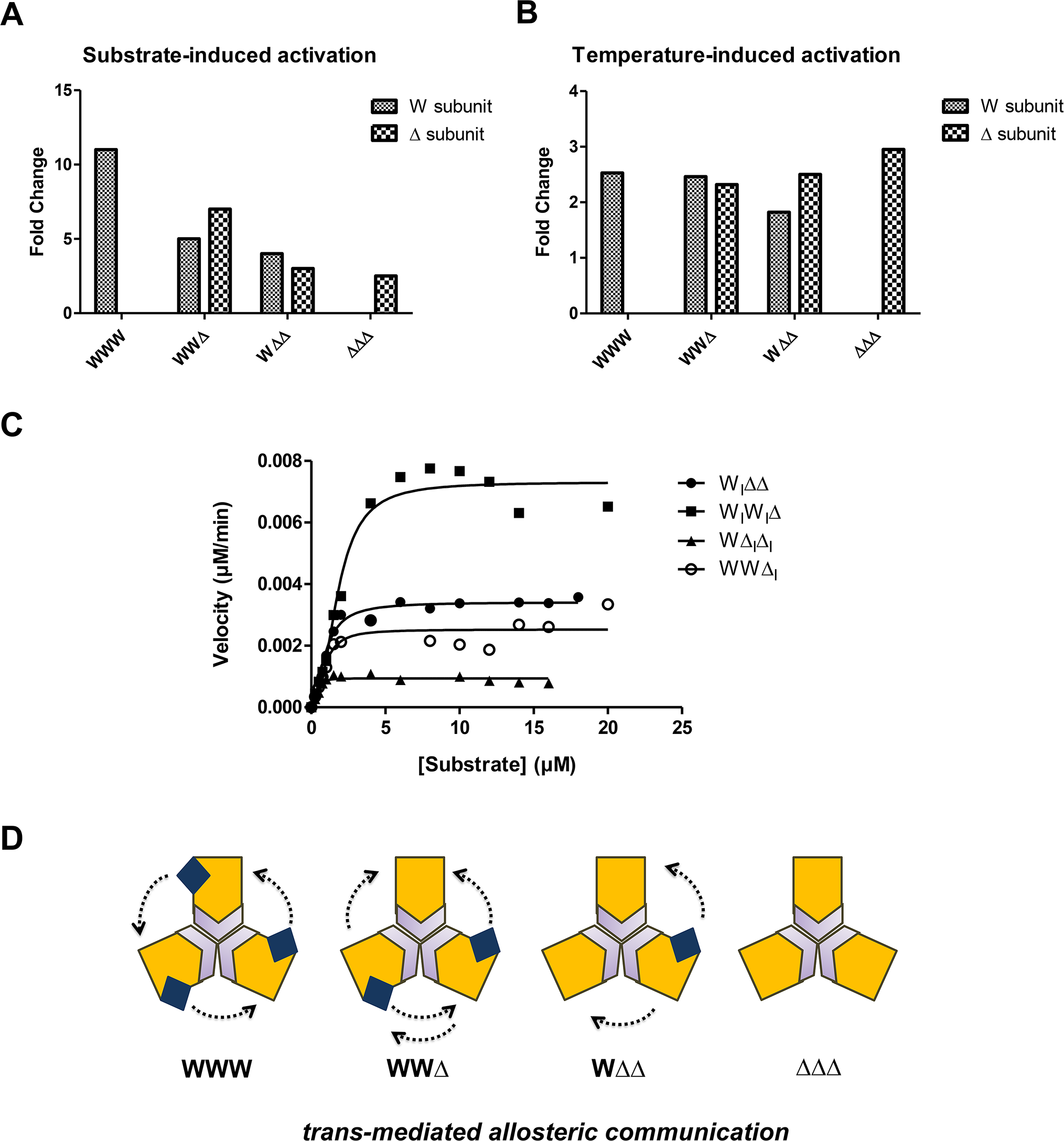
*Trans*-mediated allosteric communication under different activation modes. **(A)** Active-site modification assay performed in presence of substrate. Each variant (2 µM) was reacted with TAMRA-FP (20 µM) in absence and presence of β-casein at 37 °C for 30 min. The reaction mixtures were run on SDS-PAGE, and fluorescence scan of the complete gel was taken using ChemiDoc™ MP Imaging System. The fluorescence intensities for the W and Δ subunits in each variant were quantified using the Image Lab^TM^ software (version 6.0.0 build 25). The values obtained for each subunit from multiple independent experiments were averaged and the fold changes between the tests with respect to their corresponding controls were plotted using Graphpad Prism software. **(B)** Active-site modification assay performed at increased temperature. Each variant (2 µM) was reacted with TAMRA-FP (20 µM) and incubated either at 37 °C (Control) or at 50 °C (Test) for 30 min. The fluorescence imaging analysis and quantification were done similarly as in (A). **(C)** Quantitative assessment of proteolytic activity of HtrA2 variants with active-site mutation. Initial velocities V_0_ of HtrA2 and its variants were calculated using FITC labeled–β casein as the substrate. The solid lines are the nonlinear least squares fit of the data to the Hill form of the Michaelis–Menten equation: Velocity = V_max_/[1 + (K_0.5_/[substrate])^n^] using Graphpad Prism software. **(D)** Model representing the *trans*-mediated allosteric communication between the adjacent subunits of HtrA2 trimeric ensemble.

Taken together, our studies affirm occurrence of *trans*-mediated allosteric communication that is extremely essential for altering the active-site environment of the trimeric ensemble.

#### Temperature-based activation

For understanding heat-induced PDZ-protease plasticity, similar enzyme-modification assays were carried out by incubating each variant at 50 °C, a temperature at which both WWW and ΔΔΔ are structurally stable with enhanced proteolytic activity [17] (Figs. S3B and S3D). Interestingly, although W subunit in WWW showed maximum modification, the difference in activity fold-change with respect to the basal state in WWW, WWΔ and WΔΔ were quite comparable (Fig. 4B and Table S3B). Strikingly, the modification of Δ subunit was found to be inversely proportional to the number of PDZs present in each variant, with ΔΔΔ exhibiting maximum change in the heat-induced activation. This apparent behavioral anomaly might be attributed to the difference in structural dynamism of the protease in the two distinct activation modes. The heat-induced activation is primarily dependent on increased plasticity of PDZs in the trimer due to increase in the overall kinetic energy of the ensemble. This might lead to enhanced movement at the regulatory loops and the PDZ-protease interface. With no substrate-mediated conformational changes, the mechanism plausibly relies predominantly on the spatial dynamism of the PDZs. Thus, while WWW and WWΔ experience substantial steric hindrance from the adjoining PDZs, a single PDZ of WΔΔ remains relatively free to modify the adjacent Δ subunits. In ΔΔΔ, however, increase in activity might correlate with opening up of the active-site as a function of heat, which corroborates well with MDS analysis (Fig. 3H) and a previous FRET-based study [17]. Overall, these results hint towards a *trans*-mediated allosteric communication within the heat-activated HtrA2 protease.

### *Trans* PDZ-protease crosstalk predominates the allosteric communication in HtrA2

To further validate the intermolecular communication, we generated heterotrimeric HtrA2 variants that contained inactivated protease domain in parallel with the PDZ(s). To develop such variants, we replaced active-site S306 residue(s) with alanine (denoted as ‘I’) in WWW and ΔΔΔ constructs. Using proper subunit combinations, we generated four different heterotrimeric combinations - W_I_W_I_Δ, W_I_ΔΔ, WWΔ_I_ and WΔ_I_Δ_I_. Comparison of the enzymatic parameters of these engineered proteases with their corresponding active counterparts (WWΔ and WΔΔ) showed reduction in the β-casein substrate cleavage rate. Interestingly, despite presence of two active-sites in W_I_ΔΔ, it cleaved ∼90% of the substrate in 30 min; while W_I_W_I_Δ (with a single active-site in Δ) showed complete cleavage within the same time span (Fig. S4A). Although W_I_W_I_Δ exhibits similar enzymatic parameters as WWΔ (Fig. 4C), there is a marked 1.5-fold decrease in V_max_ and k_cat_ values of W_I_ΔΔ with respect to WΔΔ (Table 1). The catalytic efficiencies of W_I_W_I_Δ and W_I_ΔΔ are slightly lesser than their active counterparts. Interestingly, the catalytic efficiency of W_I_W_I_Δ is ∼1.5-fold greater than that of WWΔ_I_, thus strongly ascertaining the existence of *trans*-mediated allosteric communication via the PDZs. Moreover, WWΔ_I_ exhibited similar catalytic efficiency as that of WΔ_I_Δ_I_, despite having two active-sites in subunits with intact PDZs. Thus, the number of active-sites within the trimeric ensemble is not the primary defining factor for the total residual activity. This is conceivably because loss of active-sites is compensated by PDZ-mediated conformational changes that might be necessary for a well-formed oxyanion hole and subsequent substrate catalysis. In both, substrate- and heat-induced activity-based chemical profiling assays, the Δ subunit in W_I_W_I_Δ showed slightly greater modification by TAMRA than W_I_ΔΔ. Thus, similar observations from two distinct and independent studies strongly hint towards adoption of a unique compensatory mechanism, where the PDZ domain(s) strive(s) to indemnify the loss of activity of the attached subunit(s). Taken together, these results evidently prove the existence of a strong *trans*-mediated allosteric communication via a coordinated PDZ movement within the multimeric protein ensemble (Fig. 4D).

### Removal of PDZ affects formation of secondary structural elements

To delineate the role of PDZ deletion on the overall structural integrity of HtrA2 trimeric ensemble, we determined the crystallographic structure of ΔΔΔ (S306A) variant (PDB ID: 7VGE) at 4 Å resolution (Table S4). Following the trend of relatively lower resolution structures of other available ΔPDZ HtrAs (∼3 Å), it can be directed to the inherent dynamism and reduced compactness of the protease sans PDZ rather than any experimental deficit. Interestingly, majority of the ΔPDZ structures in HtrA family displayed presence of more than one trimer molecule in the crystal asymmetric unit, probably to increase stabilization and encourage crystal packing. The structural asymmetric unit of ΔΔΔ HtrA2 also contains six subunits assembled into two discrete trimers (chains - ABC and DEF) that stack against each other in a non-superimposable tail-to-tail arrangement (Fig. 5A), resulting in occlusion of the funnel-shaped surface with a buried surface area of 12610 Å^2^. The individual Δ subunits of each trimer are symmetrically packed around a three-fold molecular axis, with each subunit exhibiting slight variations in the catalytic and loop regions. Although overall the trimers are comparable to that of full-length HtrA2 (PDB ID: 1LCY) with main-chain backbone RMSD of <0.45 Å and both missing the sensory L3 loop, each Δ subunit displayed deviations near the activation domain and regulatory loops. Fortunately, despite relatively low resolution, the activation domain loops viz. LA (residues 170-174) in Chain D and L2 (residues 323-329) in Chains A and C could be successfully constructed.

**Figure 5:**
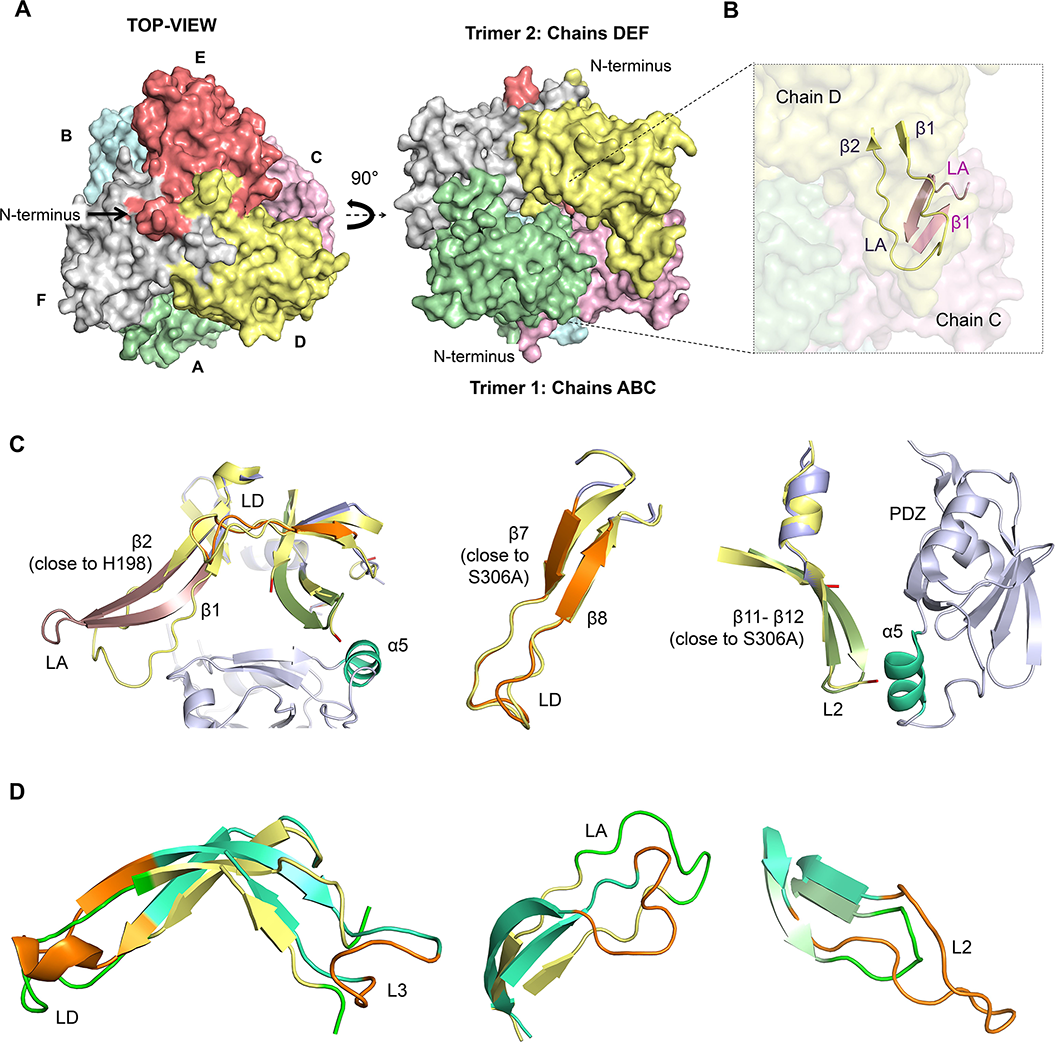
Structural features of inactive HtrA2 ΔΔΔ variant (PDB ID: 7VGE). **(A)** Surface representation of the two ΔΔΔ trimers (ABC and DEF) obtained in the asymmetric unit. Left panel: Top-view of the association, with the N-terminus of one trimer (DEF) pointing outwards. Right panel: Side-view of the association, with the two trimers having their respective N-terminus pointing in opposite directions. Chains A, B, C, D, E and F are represented in pale green, pale blue, pink, pale yellow, mauve and grey respectively. **(B)** Cross-section view showing the arrangement of LA loop and the corresponding β1-β2 between two alternate chains. In the absence of C-terminal PDZ domain, a single trimer (ABC) appears to stabilize itself by associating with another similar trimer (DEF) via the LA loop. **(C)** Superposition of full-length HtrA2 (PDB ID: 1LCY – light blue) with ΔΔΔ (pale yellow). Left panel: Orientation of LA loop of 1LCY (salmon) and ΔΔΔ (Chain D) shows the absence of secondary structure for β1-β2 in ΔΔΔ. Middle panel: β7-β8 followed by LD loop (orange in 1LCY) forms an extended secondary structure in ΔΔΔ. Right panel: Absence of α5 helix (green cyan) of the PDZ domain might result in a partial loss of secondary structure of β11-β12, and absence of L2 loop (deep olive in 1LCY) in ΔΔΔ variant. **(D)** Superposition of DegS^ΔPDZ^ (PDB ID: 3LGI – green cyan) with ΔΔΔ (Chain D – pale yellow; Chain A – pale green). Conformational changes in 3LGI and ΔΔΔ are depicted in orange and green, respectively. Left panel: The LD loop shows minor conformational changes, while the L3 loop is completely missing in ΔΔΔ (chain D). Middle panel: The LA loop displays significant difference in its orientation in 3LGI and ΔΔΔ (chain D). Right panel: Differences in orientation of L2 loop in 3LGI and ΔΔΔ (chain A). All the images were generated using PyMOL Version 1.3 (Schrodinger, LLC).

#### Inter-trimer interactions and interface analyses

Interface analysis pertaining to trimer-trimer interactions revealed that LA loop from each subunit of the upper trimer (ABC) stabilized the corresponding LA loop from the alternate subunit of the bottom trimer (DEF) (Figs. 6B and S4A). This might be the plausible reason for the two trimers to co-exist in a single asymmetric unit during crystal packing. Since PDZs are spatially close to the corresponding LA loops in the full-length structure, we speculate that absence of PDZ (especially β15-α6 region) affects the stabilization of LA loop in ΔΔΔ. The association of two trimers may further assist in the stabilization of highly dynamic LA loop (Fig. 5B). In addition, the inter-trimer interactions in ΔΔΔ are mostly non-bonded contacts between residues of LA, LD and L2 loop, with L2 residues (324-327) forming major non-bonded interactions in the inner core of the trimer association. Thus, in the absence of PDZs, LA, LD and L2 loops might play critical role in stabilizing the trimeric ensemble of ΔΔΔ HtrA2.

**Figure 6:**
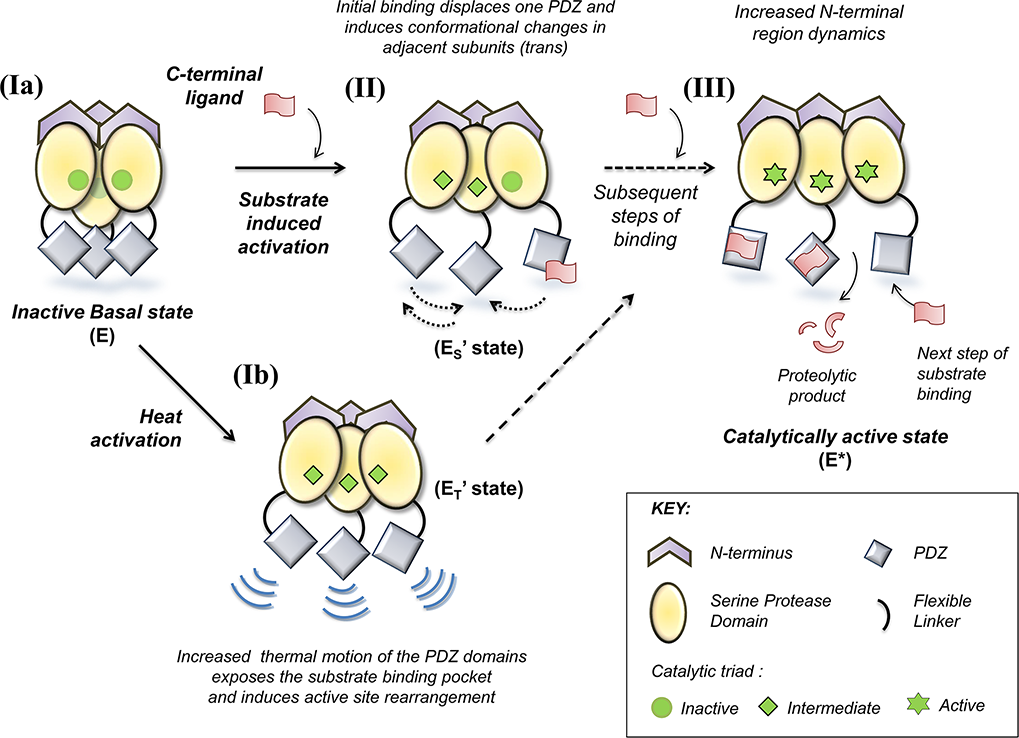
Model depicting different modes of HtrA2 activation. HtrA2 in the basal state exhibits very little activity, as the PDZs guard the entrance of the pyramidal ensemble and restrict the substrate entry (step Ia). Initial binding of the ligand at the exposed allosteric pocket of the PDZ of the original inactive state (E) induces subsequent conformational changes at the PDZ-protease domain interface (E_S_’) (step II). On the contrary, in case of activation via heat shock (step Ib), the exposed PDZs of the mature inactive protease experience increased thermal motion and induce favorable conformational changes around the active-sites in the adjacent subunits (E_T_’). The primary binding or activation event leads to displacement of a single PDZ, which further commences the synergistic coordination and formation of new interaction networks between the PDZ and protease domains of adjacent subunits (E_S_’ or E_T_’), subsequently increasing the affinity of other subunits towards the substrate by opening up the ensemble (step III) including the N-terminal region as well as rearranging the catalytic triad orientation (E*) for enhanced catalysis.

#### Structural comparisons and subunit interaction analysis

Notably, LA loop in chain D of ΔΔΔ assumed a greatly disordered conformation in contrast to the extended anti-parallel β sheet (β1and β2) observed in 1LCY (Fig. 5C). Alternatively, β7 gained an extended β-sheet conformation with respect to the positioning of the regulatory loop LD in ΔΔΔ (Figs. 6C and S4C). The β11-β12 region of L2 loop in HtrA2 (M323, V325, I329, F331) has been reported to interact with α5 and β14 of PDZ (M365, L367, I373 and L377) [15]. The abrogation of this interaction in ΔΔΔ resulted in reduced electron density for L2 loop residues in the Δ subunits (except chains A and C) (Figs. 6C and S4B). The deletion might have adversely affected the protein stability by abrogating a series of intramolecular PDZ-protease hydrophobic contacts that subsequently culminate in a disordered L2 loop with enhanced plasticity. Although the intermolecular salt bridges in most Δ subunits are similar to the wildtype structure (1LCY), PDZ deletion also affected the number of intramolecular salt bridges in ΔΔΔ. However, one of the conserved salt bridges between D190 (OD2) and R233 (NH2) remained intact despite PDZ deletion, thus underscoring its importance in maintaining intramolecular integrity of HtrA2 subunits. The structure depicts that all the disordered regulatory loops i.e., LA, L2 and LD are poised close to the comparatively more exposed catalytic triad in ΔΔΔ, than in wildtype (1LCY). Interestingly, deletion of PDZ also breaks the inter-subunit hydrogen bond between Y147—Q146 at the N-terminus. Thus, PDZ deletion significantly influences the N-terminal conformation of each subunit of HtrA2 trimeric ensemble. Furthermore, structural comparison of ΔΔΔ with relatively active bacterial counterpart, DegS^ΔPDZ^ (PDB ID: 3LGI) demonstrated several significant differences in the regulatory loop movements (RMSD <2.9 Å). Although there were no prominent changes in the orientation of the catalytic triad residues, major differences existed in the orientation of loops LA, LD, L2, L3 and their associated secondary structures (Fig. 5D).

Taken together, secondary structural perturbations around activation domain and the regulatory loops seem to play a significant role for decline in ΔΔΔ enzyme activity. These structural analyses are in concurrence with our functional assays, thereby highlighting the importance of PDZs in restructuring the orientation of regulatory loops near ΔΔΔ catalytic triad.

## DISCUSSION

Unlike its homologs, regulation of HtrA2 is multi-layered, plausibly to safeguard untimely proteolysis of its wide array of substrates. Despite numerous efforts, both discrete and collective roles of HtrA2 subunits in promoting the allosteric coordination within the trimeric ensemble remained abstruse till date. Although human HtrAs contains a single PDZ (unlike bacteria), the varying PDZ-mediated regulation suggests an evolutionary diversification of the protease structure across and within species to cater to distinct functional requirements. While various studies emphasize on HtrA2 allostery from a global perspective, the minutiae of the intricate intra- and inter-subunit networking via the regulatory PDZs remain unperceived. Furthermore, a recently published parallel study [28, 29] depicting mathematical model of interdomain interaction raises several questions due to its inability to portray the biological relevance behind experimental interpretations. In this current study, we demonstrate PDZ-mediated synergistic coordination among different domains and subunits of the protease from a biological perspective. Using heterotrimeric HtrA2 variants with varying number of PDZs, we elucidated how individually and/or in unison this domain regulates protease activation. The decline in substrate-induced proteolytic activity of the heterotrimers (WWΔ and WΔΔ) strongly suggests that the contribution of each subunit towards the total residual enzyme activity is defined by the structural identity of both the neighboring subunits, and that the subunits together exist in a cooperatively-shared energy landscape. Interestingly, in contrast to DegS, where the basal activity is inversely proportional to the number of PDZs, our findings demonstrate positive regulation of HtrA2 enzymatic functions by this regulatory domain. Moreover, our work reiterates that sequential removal of PDZs adversely affects thermal stability [17] thus highlighting its role in maintaining structural integrity.

The structural intricacy in HtrA family has often complicated the understanding of their mechanisms and hence, presence of multiple contradicting reports on their allosteric modulation is not unanticipated. For example, similar to opposing theories on DegS activation [10, 30–32], Chaganti *et. al.,* [17] challenged HtrA2 structural studies [15] and proposed a *trans* PDZ-protease crosstalk-mediated allosteric regulation of HtrA2. The use of activator-cum-substrate β-casein for the chemical profiling studies assisted in capturing the concerted and synergistic distal binding as well as cleavage mechanism from a more physiological viewpoint. The TAMRA-based active-site modification studies strongly imply the existence of a concerted *trans*-mediated pathway, wherein absence of one or two PDZs in heterotrimers (mimicking ligand-bound PDZs in cellular milieu) provided an opportunity to the adjacent subunit with intact PDZ(s) to have greater access to the PDZ-lacking subunit, thus allosterically modifying its active-site and preparing it for subsequent substrate binding. The very crowded trimeric ensemble in the apo-HtrA2 does not provide much room for such dynamic behavior and hence, might be energetically unfavorable for catalysis. This string of riveting information stimulates a precisely controlled concerted activation mechanism, which might be critical towards participating in several pathways involving a wide repertoire of substrates Similar TAMRA-based studies with heterotrimers lacking one or more active-site serine, along with quantitative enzyme kinetic analyses decidedly substantiate our claim. We have also demonstrated a concerted intermolecular PDZ-protease crosstalk in case of temperature-induced activation of HtrA2. However, the observed distinct behavior of the heat-based activation might be attributed to the overall dynamism accrued by the PDZs due to rise in temperature. With removal of the PDZ inhibition (ΔΔΔ), activation via temperature might cause certain favorable rearrangements at the N-terminal region [17] and around the active-site such that it compensates for the lost PDZs by opening up the ensemble.

To resolve the ambiguities pertaining to distinct activation pattern in HtrA2 and ΔΔΔ, we reported the structure of catalytically inactive (S306A) version of ΔΔΔ. As the ΔΔΔ and wild type (PDB ID: 1LCY) structures share similar orientation of catalytic triad residues, we believe that increase in temperature or the binding of specific ligands (activator or substrate) greatly influences the PDZ dynamics and subsequently the active-site reactivity by bringing about suitable regulatory loop rearrangements with respect to the oxyanion hole. For DegS, which is a more open ensemble, and where PDZs act as inhibitory modules, deletion of PDZs do not greatly affect the catalysis of this variant [33–35]. On the contrary, PDZs in HtrA2 seem to play a greater role in structural stabilization by assisting secondary structure formation in protease domains, especially around the regulatory loops that together form the activation domain, which is lost in the ΔΔΔ variant. Thus, in the absence of any form of activation, PDZs maintain the pyramidal architecture in a closed conformation by stabilizing intramolecular PDZ-protease domain interactions. Absence of inter- and intra-subunit interactions in ΔΔΔ, which could have otherwise favorably assisted establishment of the required transitions (L3 to LD* loop via L1*/L2* (asterisk denotes adjacent subunits)) to the catalytically competent state [16], and also energetically stabilized the protease, reiterates the indispensability of these regulatory domains in upholding the overall protease structural integrity and functions.

Overall, our observations propound a working model for HtrA2 activation, where the initial step of ligand binding or heat activation shifts the intrinsic equilibrium from an inactive state (E) to a higher affinity conformation (E’) that eventually promotes a more catalytically competent state (E*) through a synergistic *trans* PDZ-protease coordination, as elaborated in Figure 6. Briefly, ligand binding at the exposed substrate binding pocket of PDZ brings about subsequent conformational changes at the PDZ-protease interface that are different from the original inactive state (E). The primary binding leads to displacement of a single PDZ (E_S_’ state), which further commences a synergistic coordination and formation of new interaction networks between the PDZ and protease domains of adjacent subunits, subsequently increasing the affinity of other subunits by opening up the ensemble as well as reorienting the catalytic triad for enhanced catalysis. On the other hand, in temperature-mediated activation, (Fig. 6) the increase in temperature promotes inter-domain PDZ movement in a way that not only reduces the steric crowd but also leads to subtle distal conformational changes conducive toward competent active-site formation. Thus, under cellular conditions, stress-related temperature change might induce initial opening of PDZs, in turn making the catalytic pocket more accessible for substrate binding and proteolysis. Furthermore, this temperature-induced reordering of the active-site (E_T_’ state) might be particularly important in scenarios where the protease-domain defines initial substrate binding and its subsequent cleavage (unpublished data). Thus, this model describes an energy landscape that might be allosterically tuned in multiple ways to help the protease to conform to different cellular requirements for optimal biological leverage. This kind of dynamic perpetuity, observed in a multimeric protease ensemble through a discrete and sequential inter-domain coupling, allows a robust coalescence of regulatory strategies amongst heterogeneous sub-ensembles that successfully culminate in an amenable functional conformation [36].

In conclusion, this manuscript unequivocally describes the *trans*-mediated allosteric mechanism of HtrA2 in two different activation modes using a unique retrospective approach through dissecting and rebuilding the protein structure. This study might open up new avenues for devising tailor-made strategies toward modulating diverse functions of HtrA2 with desired characteristics for therapeutic benefit. Most importantly, delineation of this multidimensional allosteric model system that amalgamates both activation and stabilization of the macromolecular ensemble through an intricate intermolecular crosstalk for transmission of information within and outside cellular pathways provides an advanced understanding of allostery in general from a more quantitative perspective.

## DATA AVAILABILITY

The newly solved crystal structure reported in this publication is available in RCSB PDB database (http://www.wwpdb.org) under accession number 7VGE (PDB DOI: https://doi.org/10.2210/pdb7VGE/pdb). All the other data are available in the main text or the supplementary materials. Additional data related to this paper may be requested from the authors.

## ACKNOWLEDGEMENTS

We thank LK. Chaganti, N. Singh and A. Kalarikkal for their intellectual inputs and helpful suggestions related to experimentation. We are grateful to the XRD facility IIT Bombay for initial crystallization screening and crystallographic data collection. This work is supported by the Department of Biotechnology (DBT), Govt. of India (grant number BT/HRD/NWBA/37/01/2015) and intramural research grant received from ACTREC-TMC, India (IEC project no. 162).

## AUTHOR CONTRIBUTIONS

Conceptualization: ALP and KB

Methodology: ALP, VM and PB (structure), SD (*in-silico*)

Investigation: ALP, VM and PB (structure), SD (*in-silico*)

Visualization: ALP, VM, SD

Supervision: ALP, PB, KB

Writing—original draft: ALP, SD (*in-silico*) and KB

Writing—review & editing: ALP, VM, SD, PB and KB

## STAR METHODS

### Recombinant protein expression and purification

The mature form of HtrA2 full length protein, comprising residues 134–458, is considered as Wildtype (WT or WWW) and the PDZ lacking HtrA2 variant containing the N-terminus-SPD domain (residues 134-210) is considered as N-SPD or ΔΔΔ. These proteins are cloned and expressed in pET-20b (Addgene, Cambridge, MA, USA) and modified pMAL-c5E vectors (New England Biolabs, Ipswich, MA, USA) respectively, as previously described [17]. The latter had a TEV (Tobacco Etch Virus) protease cleavage site introduced for removal of the fusion tag. An additional FLAG octapeptide (DYKDDDDK) tag at the N-terminal end in ΔΔΔ construct and single amino acid substitution of S306 residue to Alanine in both the constructs (WWW and ΔΔΔ) was performed using site-directed mutagenesis (Stratagene, Austin, TX, USA). The insertion and point mutations were verified by automated DNA sequencing. Recombinant proteins were expressed in *E. coli* BL21 (DE3) pLysS and Rosetta (DE3) pLysS strains (Novagen, Billerica, MA, USA). These cells were grown at 37 °C until the optical density at 600 nm reached ∼0.6 AU and the protein expression was then induced by adding isopropyl β-D-1-thiogalactopyranoside (IPTG) to a final concentration of 0.3 mM. The cells were further cultured at 18 °C for 18 h post induction and then harvested for protein production. Protein variants with a C-terminal Histidine tag were purified by affinity chromatography using Ni-NTA resin (Nucleo-pore, Genetix Biotech Asia Pvt. Ltd), while those with N-terminal maltose binding protein (MBP) fusion tag were purified using amylose resin (New England Biolabs). All the protein variants were purified in buffer A (20 mM Na_2_HPO_4_/NaH_2_PO_4_ (pH 8.0) containing 100 mM NaCl and 3% (v/v) glycerol) and the proteins obtained were >95% pure, as estimated by SDS PAGE.

### Generation and purification of heterotrimeric HtrA2 variants

For obtaining heterotrimeric variants, the use of simultaneous heat and chemical denaturation process was employed. Purified WWW and ΔΔΔ trimers were mixed in equimolar concentrations and diluted up to 15-fold in buffer A, additionally containing 6 M guanidinium chloride (pH 8.0). This reaction mixture was further subjected to heat by incubating at 50 °C for 1-2 h for efficient denaturation into monomeric subunits. The solution was further dialyzed extensively against a renaturation buffer (buffer A (pH 8.0) containing 300 mM arginine with varying content of glycerol) at 4 °C. The final step involved dialysis in buffer A (to remove arginine) and concentrating the renatured protein mixture using an Ultracel-30k centrifugal filter (Merck Millipore). For further separation of the different HtrA2 variants (homo- and hetero-trimers), this renatured mixture was bound to Ni-IDA column pre-equilibrated with buffer A. Since the variants differed in the total number of histidine residues (WWW-His_6_ and ΔΔΔ-His_3_) at their C-terminal end, a very shallow gradient of imidazole (10-250 mM) in the elution buffers was used for their separation. The impurities eluted in the flow-through fraction and up to 30 mM imidazole buffer washes, followed by WΔΔ, WWΔ and WWW eluting between 30 – 250 mM imidazole containing elution buffers. The elution fractions were loaded on SDS PAGE for analyzing the fractions containing the heterotrimers.

### Native PAGE and Western Blot

To confirm the identity of the generated HtrA2 heterotrimers, protein samples from the appropriate purification fractions were mixed with native PAGE sample buffer (62.5 mM Tris-Cl, pH 6.8, 15% glycerol, and 0.01% bromophenol blue) and loaded on 7.5% native PAGE. The gels were run at a constant voltage of 50 V for 5 h at 4 °C and were analyzed by Coomassie Blue staining. Since all the homo- and hetero-trimers also differed in the number of FLAG tag (originally introduced in ΔΔΔ construct), the unstained native PAGE gel was additionally probed with monoclonal anti-FLAG antibody by performing Western blot. The separated protein bands were transferred on nitrocellulose membrane (Millipore, Italy) using Invitrogen wet transfer mini blot module (Thermo Fisher Scientific, USA) in 1X transfer buffer (25 mM Tris, 192 mM glycine, 20% (v/v) methanol, 0.025–0.1% SDS, pH 8.3) at a constant voltage of 15 V for 60-90 min. The membrane was then blocked with 5% bovine serum albumin (BSA) prepared in 1X TBST buffer (Tris-buffered saline (TBS) containing 0.1% Tween-20) for 1h, followed by overnight incubation with 1:2000 dilution of anti-FLAG antibody (Sigma Aldrich) at 4 °C. The membrane was further incubated at room temperature with 1:5000 dilution of goat antimouse IgG HRPO (Sigma Aldrich) secondary antibody for 1 h. After each incubation period mentioned above, the membrane was washed three to six times with 1X TBST solution. Antibody dilutions were prepared in TBS containing 0.01% Tween-20 and 1% BSA. Bands were developed using an enhanced chemiluminescence kit (Thermo Fischer, USA).

### Protease assays

Protease activity of different HtrA2 variants (homo- and hetero-trimers) was semi-quantitatively determined using the generic serine protease substrate β-casein (Sigma, St. Louis, MO, U.S.A.). For each 30 µl reaction, 2 µg of respective protein variant was incubated with 6 µg of β-casein in buffer A at 37 °C for a course of different time points (0-90 min). The reaction was stopped with a 5X SDS sample loading buffer, followed by boiling for 10 min. The cleavage products were resolved on 15% SDS PAGE and the results were analyzed by Coomassie staining. Protein bands were quantitated using *GelQuant.NET* software. The data was normalized with respect to the uncleaved substrate control and was plotted using the GraphPad Prism 5 (GraphPad Software, San Diego, CA, USA). For all quantitative studies, fluorescein isothiocyanate (FITC) β-casein (Sigma, St. Louis, MO, U.S.A.) was used as the substrate. For each 100 µl reaction mixture, 2 µM protein was incubated with increasing concentrations (0-20 µM) of FITC β-casein at 37 °C in buffer A. The rate of proteolytic cleavage was monitored in Cytation^TM^ 5 multi-mode plate reader (BioTek Instruments, Inc.) using an excitation wavelength of 485 nm, followed by emission at 535 nm. For each HtrA2 variant, the initial velocities V_0_ (μM/min) were calculated using linear regression analysis. The steady-state kinetic parameters were calculated from the reaction rates by fitting the data to the Hill form of Michaelis–Menten equation, Velocity = V_max_/[1 + (K_0.5_/[substrate])^n^] (where V_max_ represents the maximum velocity and K_0.5_ is the substrate concentration at half maximal velocity) in GraphPad Prism version 5.01 for Windows (GraphPad Software, San Diego, California USA). All the experiments were performed independently in triplicates and the mean SEM values were reported.

### Activation assay using chemical probe

For trapping the active-site reactivity, chemical profiling of the enzyme activity was performed using ActivX TAMRA-FP Serine Hydrolase Probe (Thermo Scientific, USA). 0.1mM stock of TAMRA-FP was prepared by dissolving the commercially available probe in anhydrous dimethyl sulfoxide, according to the manufacturer’s protocol. For each reaction mixture, a final concentration of 2 µM probe was added and the reactions were incubated for 30 min. For substrate-based and temperature-induced activation, the incubation temperatures were set to 37°C and 50 °C respectively. The reactions were terminated by adding 5X SDS sample loading buffer, followed by boiling for 10 min. The labeled proteins were resolved on 15% SDS PAGE and the results were analyzed using ChemiDoc™ MP Imaging System (Bio-Rad laboratories, Inc.) with Cy®3 filters. For better quantitative analysis, the fluorescence intensities of each subunit band were measured using Image Lab^TM^-version 6.0.0 build 25 (2017, Bio-Rad laboratories, Inc.) and the fold change between the intensities of the subunits in the test samples with respect to their corresponding controls were then plotted for graphical representation.

### *In silico* preparation of the heterotrimeric models of HtrA2

For preparation of *in silico* heterotrimeric models, three-dimensional (3D) crystal structure of HtrA2 (PDB ID: 5M3N) was retrieved from Protein Data Bank [37]. The 3D structure lacked crystal information for few loop regions-an N-terminal loop (^1^AVPSPPPA^8^), a loop in the SPD region (^280^RPARDLGLPQTNV^292^) and the linker region (^344^RGEKKNSSSGISGS^357^) [37]. Hence, these missing regions were modeled using *ab initio* mode of loop filling and protein preparation programs in Prime (Prime, Schrödinger, LLC, New York, NY, 2020) [38]. Loop filling was done on the basis of permissive dihedral angle values for different residues, followed by repetitive rounds of sample loop clustering, optimizing the side-chain, and energy minimization of the loops [38]. The resultant loop-filled wildtype HtrA2 monomer was energy minimized using Desmond (Desmond, Schrödinger, LLC, New York, NY, 2020). Using the crystal symmetry data of HtrA2 monomer [37], the trimeric ensemble of HtrA2 wildtype (WWW) was generated, which was further modified by deletion of the linker and PDZ domain in subsequent chains, viz, WWΔ (with linker and PDZ deleted in chain C), WΔΔ (with linker and PDZ deleted in chain B and C), and, ΔΔΔ (with linker and PDZ deleted in all the three chains). These models were then subjected to docking using a β-casein peptide denoted by GPFPIIV (a generic substrate). Docking was performed using Bioluminate software (Bioluminate, Schrödinger, LLC, New York, NY, 2020) and scored using MM-GBSA (Molecular Mechanics using Generalized Born and Surface Area continuum) scoring method [39]. The top five docked complexes for β-casein-bound HtrA2 variants were analysed for the identification of interacting surface residues using PDBSum online server [40].

### Molecular dynamic simulation (MDS) analysis of bound and unbound HtrA2 variants

Both peptide-bound and unbound forms of WWW, WWΔ, WΔΔ and ΔΔΔ were subjected to MDS using Desmond (Desmond, Schrödinger, LLC, New York, NY, 2020) where OPLS3e (Optimized Potentials for Liquid Simulations Version 3e) force field was used to generate topology and parameter files [41, 42]. Each protein structure was surrounded by a cubic box of TIP3P water molecules with the nearest distance from the protein to the box boundary being no more than 10 Å [43]. The generated systems were subsequently neutralized (net charge was brought to zero) by adding adequate number of positive (Na^+^) ions. Each system underwent one round of steepest-descent minimization, followed by one round of conjugated gradient for 5000 picosecond (ps) [44]. The systems were then equilibrated in NVT (constant number of particles, volume, and temperature) and NPT (constant number of particles, pressure, and temperature) ensembles with two sets of restrained NVT (for 24 ps and 2000 ps respectively) and one set of restrained (for 24 ps) and unrestrained (for 5000 ps) NPT each [45]. During equilibration, LINCS (LINear Constraint Solver) constraint algorithm was used to apply position restraining force on all the atomic bonds present in the systems [46]. All the systems were then subjected to final MD simulation run for 1000 nanosecond (ns) under no-restrained NPT ensemble. For peptide-bound systems, the final production runs were performed at 300 K. However, for analyzing the temperature dependent changes in the unbound systems, the final temperature was kept at 323 K. Post-simulation these systems were further subjected to meta-dynamics analysis for 50 ns that generated free energy landscape data on the basis of the active-site conformation. All the MD simulation data analysis was carried using MD simulation analysis tools available in Desmond and Maestro platform (Desmond and Maestro, Schrödinger, LLC, New York, NY, 2020).

### Protein crystallization

To eliminate the possibility of partial protein degradation at higher concentrations required for crystallization, the inactive variant of N-SPD (ΔΔΔ-S306A) was used. The protein was purified in buffer containing 10 mM HEPES (pH 8.0), 100 mM NaCl, and 3% glycerol, in a similar way as described for active ΔΔΔ. Four commercial crystallization screens from Hampton Research (HR2-110 and HR2-112) and Molecular Dimensions (Structure Screens MD1 -01 and MD1 -02) were used for setting crystallization trials using protein sample with concentration of 15 mg/ml. The optimization screen was set using hanging drop vapor diffusion after mixing protein (∼6 mg/mL) in 1:1 ratio with mother liquor comprising 0.5 M Sodium acetate trihydrate, 2.0 M Sodium formate pH 6.0, 3% glycerol, and diffraction-quality crystals were obtained after two weeks at 16 °C.

### Data collection and structure determination

Prior to diffraction data collection, the crystals were soaked in a cryo-protectant solution (mother liquor with 20% ethylene glycol) and then flash frozen in a liquid nitrogen stream at 100 K. Diffraction data were collected by the rotation method with a Rigaku MicroMax 007HF generator equipped with R-Axis IV++ detector using CuKα X-ray radiation (1.5418 Å) at the Protein Crystallography Facility, IIT Bombay, Mumbai. Diffraction data was collected with an exposure of 10 min, at a detector distance of 370 mm. The crystal was subjected to sequential steps of annealing (of 5 s each) for a total of 15 s. The diffraction images were indexed, integrated, and scaled with XDS [47]. The intensities were converted to the structure factors with the programs F2MTZ and CAD of CCP4 [48]. The data collection statistics are presented in Table S4. The calculated Matthew’s coefficient for ΔΔΔ is 2.49 Å^3^ Da^-1^ that indicated the presence of six molecules in the asymmetric unit. The phases were obtained by molecular replacement with the program PHASER [49] using the coordinates of the N-terminal region and protease domain of the previously reported structure (PDB ID: ILCY). Further rounds of manual model building and electron density interpretation was performed in COOT [50] and refinement was executed with REFMAC5 [51]. The first few N-terminal residues in an asymmetric unit and the flexible loops of certain regions could not be modeled due to the lack of proper electron density. The final refinement statistics and validation parameters obtained with MolProbity [52] of the refined structure are summarized in Table S4. All structure-related figures were generated with PyMOL (https://www.pymol.org/).

## Supplemental Information

**Table S1:**
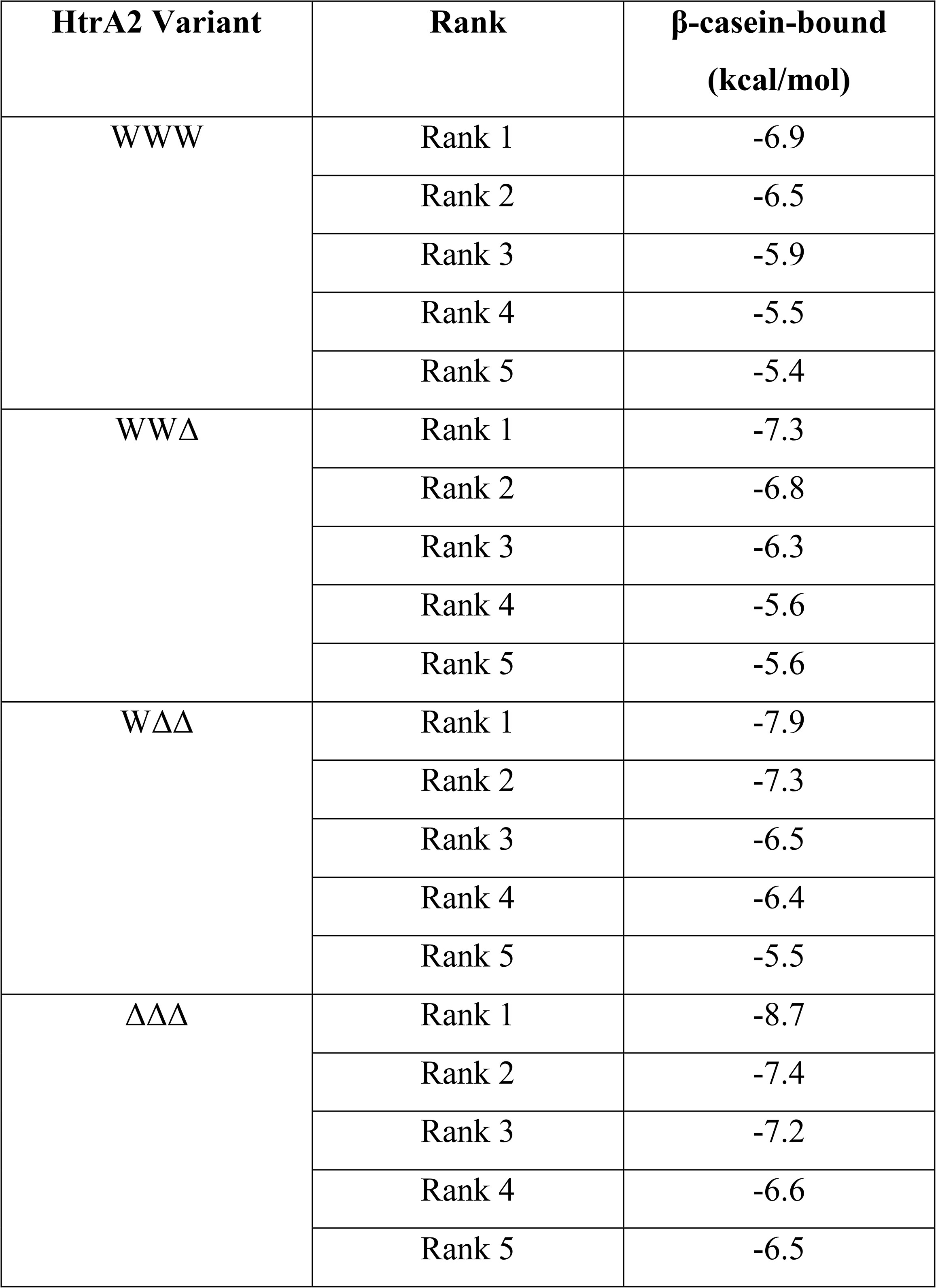
Top 5 docking scores generated for β-casein-bound WWW, WWΔ, WΔΔ and ΔΔΔ.

| HtrA2 Variant | Rank | β-casein-bound<br>(kcal/mol) |
| --- | --- | --- |
| WWW | Rank 1 | -6.9 |
|  | Rank 2 | -6.5 |
|  | Rank 3 | -5.9 |
|  | Rank 4 | -5.5 |
|  | Rank 5 | -5.4 |
| WWΔ | Rank 1 | -7.3 |
|  | Rank 2 | -6.8 |
|  | Rank 3 | -6.3 |
|  | Rank 4 | -5.6 |
|  | Rank 5 | -5.6 |
| W$\Delta\Delta$ | Rank 1 | -7.9 |
|  | Rank 2 | -7.3 |
|  | Rank 3 | -6.5 |
|  | Rank 4 | -6.4 |
|  | Rank 5 | -5.5 |
| $\Delta\Delta\Delta$ | Rank 1 | -8.7 |
|  | Rank 2 | -7.4 |
|  | Rank 3 | -7.2 |
|  | Rank 4 | -6.6 |
|  | Rank 5 | -6.5 |

**Table S2:** Average distance (in Å) analysis among the catalytic triad residues for HtrA2 variants. The distances were calculated over a 1000 ns time-scale for β-casein-bound and temperature induced WWW, WWΔ, WΔΔ and ΔΔΔ.

| Residue | β-casein-bound (in Å) | | | | | | | |
| --- | --- | --- | --- | --- | --- | --- | --- | --- |
| | WWW | | WWΔ | | W$\Delta\Delta$ | | $\Delta\Delta\Delta$ | |
|  | Unbound | Bound | Unbound | Bound | Unbound | Bound | Unbound | Bound |
| ChainA<br>H198 to<br>D228 | 2.9 ± 0.2 | 2.6 ± 0.1 | 2.9 ± 0.3 | 2.8 ± 0.1 | 3.0 ± 0.2 | 2.9 ± 0.3 | 2.9 ± 0.2 | 3.2 ± 0.2 |
| ChainA<br>H198 to<br>S306 | 4.1 ± 0.2 | 4.9 ± 0.1 | 4.2 ± 0.2 | 4.5 ± 0.2 | 4.2 ± 0.2 | 4.2 ± 0.1 | 4.1 ± 0.2 | 3.7 ± 0.2 |
| ChainB<br>H198 to<br>D228 | 2.9 ± 0.2 | 2.7 ± 0.2 | 2.9 ± 0.3 | 2.7 ± 0.1 | 3.0 ± 0.3 | 3.0 ± 0.1 | 2.9 ± 0.2 | 3.1 ± 0.2 |
| ChainB<br>H198 to<br>S306 | 4.1 ± 0.2 | 5.3 ± 0.2 | 4.2 ± 0.2 | 4.4 ± 0.1 | 4.2 ± 0.2 | 4.3 ± 0.2 | 4.1 ± 0.2 | 3.6 ± 0.1 |
| ChainC<br>H198 to<br>D228 | 2.9 ± 0.2 | 2.5 ± 0.2 | 2.9 ± 0.3 | 2.6 ± 0.2 | 3.0 ± 0.3 | 2.7 ± 0.3 | 2.9 ± 0.2 | 3.2 ± 0.1 |
| ChainC<br>H198 to<br>S306 | 4.1 ± 0.2 | 5.1 ± 0.2 | 4.2 ± 0.2 | 4.5 ± 0.1 | 4.2 ± 0.2 | 4.4 ± 0.1 | 4.1 ± 0.2 | 3.7 ± 0.3 |
| Residue | Temperature-induced activation (in Å) |  |  |  |  |  |  |  |
| | WWW | | WWΔ | | W$\Delta\Delta$ | | $\Delta\Delta\Delta$ | |
|  | Uninduced | Induced | Uninduced | Induced | Uninduced | Induced | Uninduced | Induced |
| ChainA<br>H198 to<br>D228 | 2.9 ± 0.2 | 2.7 ± 0.1 | 2.9 ± 0.3 | 2.8 ± 0.2 | 3.0 ± 0.3 | 2.9 ± 0.1 | 2.9 ± 0.2 | 3.4 ± 0.3 |
| ChainA<br>H198 to<br>S306 | 4.1 ± 0.2 | 5.5 ± 0.2 | 4.2 ± 0.2 | 4.4 ± 0.1 | 4.2 ± 0.2 | 4.1 ± 0.3 | 4.1 ± 0.2 | 3.5 ± 0.3 |
| **ChainB**<br>**H198 to**<br>**D228** | 2.9 ± 0.2 | 2.6 ± 0.2 | 2.9 ± 0.3 | 2.8 ± 0.1 | 3.0 ± 0.3 | 3.1 ± 0.2 | 2.9 ± 0.2 | 3.1 ± 0.1 |
| **ChainB**<br>**H198 to**<br>**S306** | 4.1 ± 0.2 | 4.8 ± 0.1 | 4.2 ± 0.2 | 4.5 ± 0.1 | 4.2 ± 0.2 | 4.0 ± 0.3 | 4.1 ± 0.2 | 3.6 ± 0.2 |
| **ChainC**<br>**H198 to**<br>**D228** | 2.9 ± 0.2 | 2.7 ± 0.1 | 2.9 ± 0.3 | 2.8 ± 0.1 | 3.0 ± 0.3 | 2.7 ± 0.3 | 2.9 ± 0.2 | 3.3 ± 0.3 |
| **ChainC**<br>**H198 to**<br>**S306** | 4.1 ± 0.2 | 5.2 ± 0.2 | 4.2 ± 0.2 | 4.4 ± 0.1 | 4.2 ± 0.2 | 4.5 ± 0.3 | 4.1 ± 0.2 | 3.8 ± 0.1 |

**Table S3:** Quantitative analyses of the active-site modification assay. The average fluorescence intensity values and the corresponding fold change for W and Δ subunit in the control and test samples for each HtrA2 variant.

**(A) Substrate-induced activation**
| **HtrA2**<br>**Variants** | **W Subunit** |  |  | **Δ Subunit** |  |  |
| --- | --- | --- | --- | --- | --- | --- |
|  | **Control\*** | **Test\*** | **Fold**<br>**change** | **Control\*** | **Test\*** | **Fold**<br>**change** |
| WWW | 4865 | 53518 | 11 | 0 | 0 | 0 |
| WWΔ | 11805 | 59415 | 5 | 7625.5 | 54542.5 | 7 |
| WΔΔ | 8629 | 35255 | 4 | 14469 | 46090 | 3 |
| ΔΔΔ | 0 | 0 | 0 | 2687 | 6648.5 | 2.5 |

**(B) Temperature-induced activation**
| **HtrA2**<br>**Variants** | **W Subunit** |  |  | **Δ Subunit** |  |  |
| --- | --- | --- | --- | --- | --- | --- |
|  | **Control\*** | **Test\*** | **Fold**<br>**change** | **Control\*** | **Test\*** | **Fold**<br>**change** |
| WWW | 2682.5 | 6803.5 | 2.53 | 0 | 0 | 0 |
| WWΔ | 5954 | 14646 | 2.46 | 4436 | 10314 | 2.32 |
| WΔΔ | 5803.5 | 10582.5 | 1.82 | 10639.5 | 26644 | 2.5 |
| ΔΔΔ | 0 | 0 | 0 | 2456.5 | 7258 | 2.95 |

\* Arbitrary Unit

**Table S4:** Crystallography data statistics for ΔΔΔ (PDB ID: 7VGE)

| **Data collection and refinement statistics** | **HtrA2 $\Delta\Delta\Delta$** |
| --- | --- |
| **Data collection statistics** |  |
| Space group | $P4_32_12$ |
| Unit cell parameters<br>$a, b, c$ (Å)<br>$\alpha, \beta, \gamma$ (°) | 82.88, 82.88, 395.31<br>90.00, 90.00, 90.00 |
| Temperature (K) | 100 |
| Wavelength (Å) | 1.5419 |
| Resolution (Å) | 40.0-4.0 (4.1-4.0) |
| $R_{\text{merge}}$ (%) | 39.8 (144.8) |
| Completeness (%) | 90.7 (90.9) |
| Mean $I/\sigma(I)$ | 4.36 (1.27) |
| Total reflections | 85456 (5452) |
| Unique reflections | 20149 (1441) |
| Redundancy | 4.24 (3.78) |
| $CC_{1/2}$ | 96.7 (25.1) |
| Wilson B factor (Å<sup>2</sup>) | 76.05 |
| No. of molecules in ASU | 6 |
| **Refinement statistics** |  |
| Resolution (Å) | 39.53-4.0 |
| Working set: no. of reflections | 11300 |
| $R_{\text{factor}}$ (%) | 26.51 |
| Test set: no. of reflections | 595 |
| $R_{\text{free}}$ (%) | 33.64 |
| Protein atoms | 8149 |
| **Isotropic average B-factor (Å)** |  |
| Protein | 116.3 |
| **Geometry statistics** |  |
| R.m.s.d. bond length (Å) | 0.006 |
| R.m.s.d. bond angle (°) | 1.5 |
| **Ramachandran plot (%)** |  |
| Most favoured region | 93.0 |
| Allowed regions | 6.7 |
| Outlier | 0.3 |

**Figure S1:**
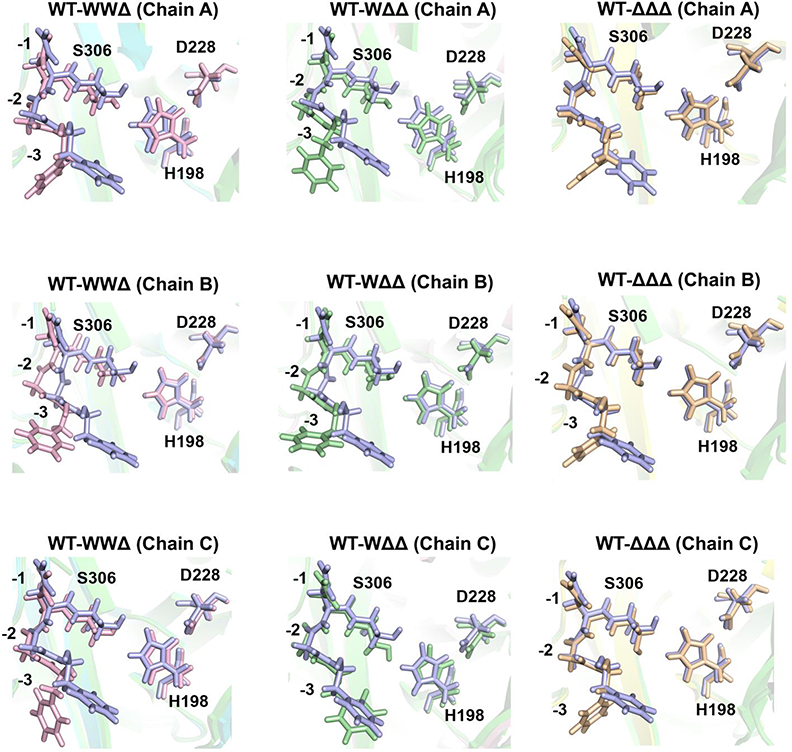
Orientation of the catalytic triad residues of HtrA2 variants. Stick diagram showing the alignment of the catalytic triad residues for heterotrimeric variants (WWΔ and WΔΔ) and ΔΔΔ (N-SPD variant) with respect to the wild type (WWW). Catalytic residues are represented as H198, D228 and S306. Oxyanion hole residues are marked as -1, -2 and -3 starting from the S306 residue which is considered as 0. WWW, WWΔ, WΔΔ and ΔΔΔ are denoted by light blue, pink, green and yellow sticks, respectively.

**Figure S2:**
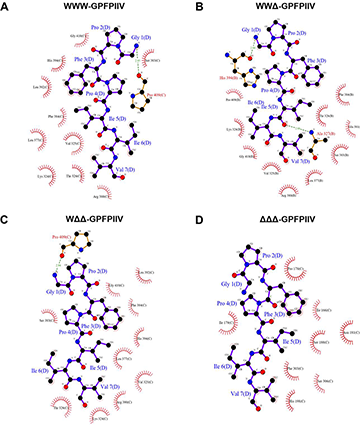
Interaction analyses of HtrA2 variants in presence of C-terminal ligand. Ligplot showing the interacting residues between the β-casein peptide (GPFPIIV) and **A**) WWW, **B**) WWΔ, **C**) WΔΔ and **D**) ΔΔΔ variants of HtrA2. Hydrogen bond interactions are denoted by green dotted lines.

**Figure S3:**
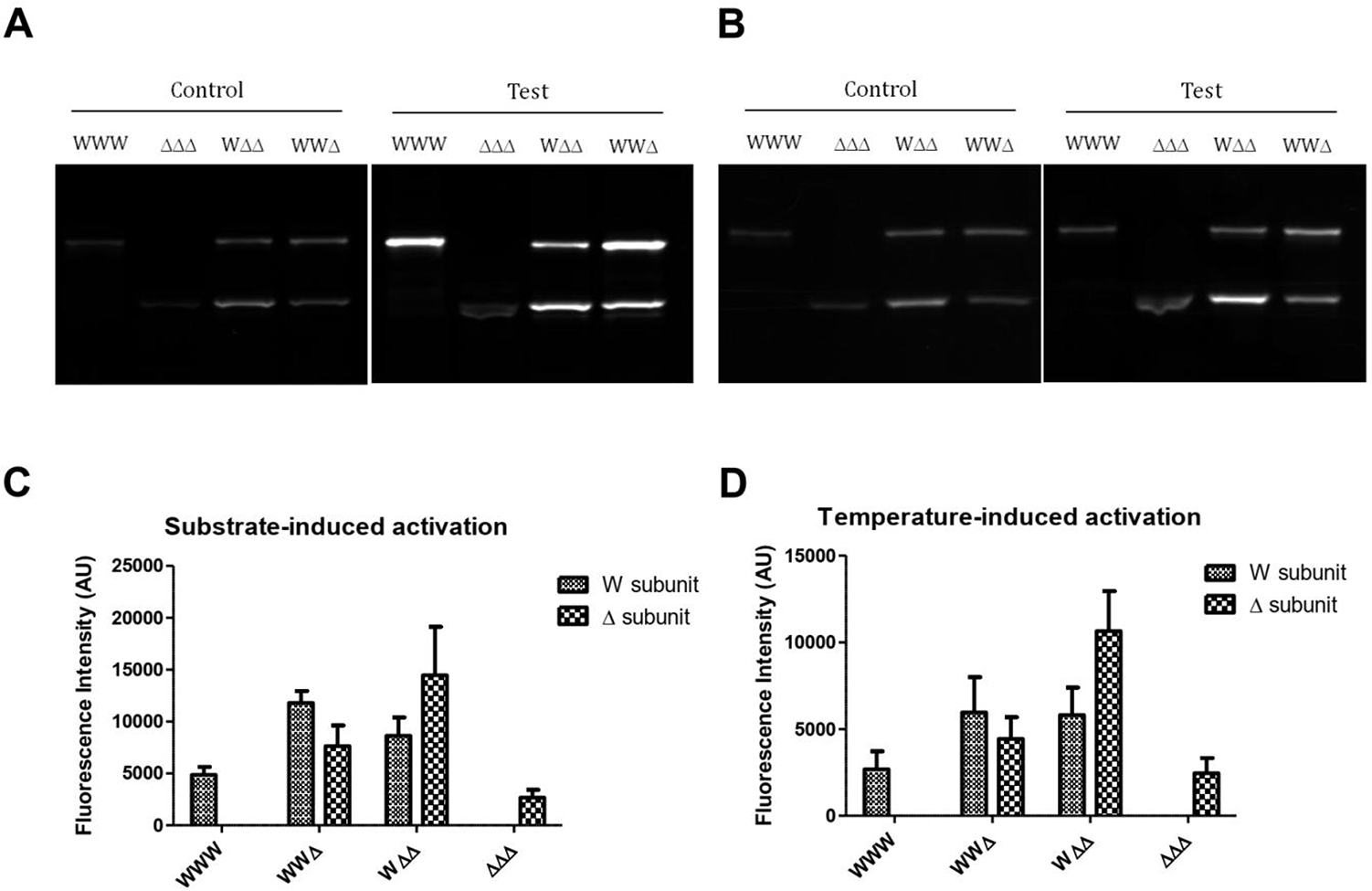
In-gel fluorescence imaging performed for active-site modification assay using TAMRA-FP. Representative images from **(A)** Substrate-induced activation **(B)** Temperature-induced activation. **(C)** The fluorescence intensities of the control samples in (A) were quantified using the Image Lab^TM^ software (version 6.0.0 build 25). The values obtained for each subunit from multiple independent experiments were plotted using Graphpad Prism software with their respective SEM. **(D)** The fluorescence intensities of the control samples in (B) were quantified and represented in a similar way as in (C).

**Figure S4:**
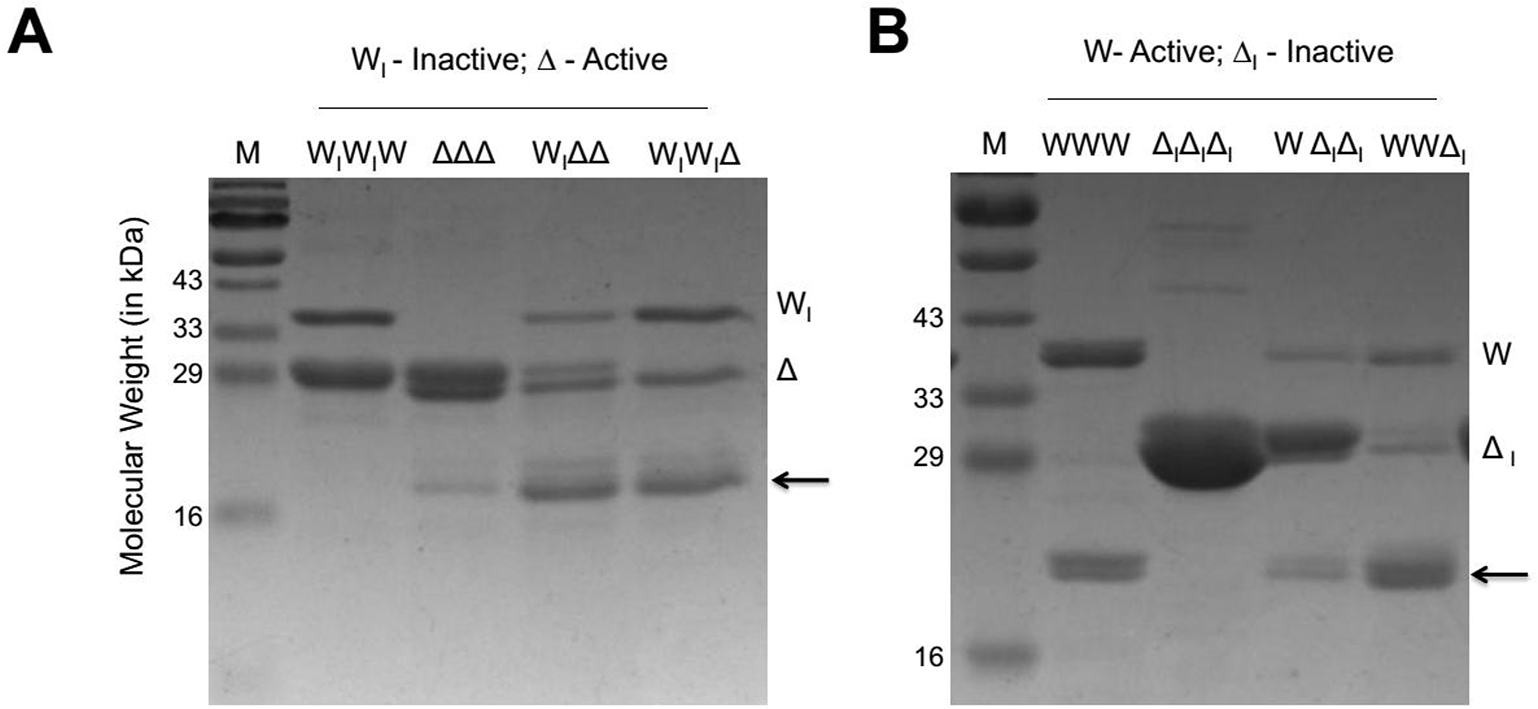
Proteolytic cleavage assays of HtrA2 variants with active-site mutation. **(A) (B)** Qualitative gel-based proteolytic cleavage of β-casein by HtrA2 variants containing active-site mutation in different subunits. 2 μg of each enzyme variant was incubated individually with 6 μg of substrate β-casein and incubated at 37 °C for 30 min. The reaction at each time point was stopped with Laemmli buffer at 100 °C. Reaction samples were resolved by 12% SDS-PAGE and the cleavage pattern was visualized with Coomassie brilliant blue staining. Arrows indicate the cleaved products. M: Protein marker.

**Figure S5:**
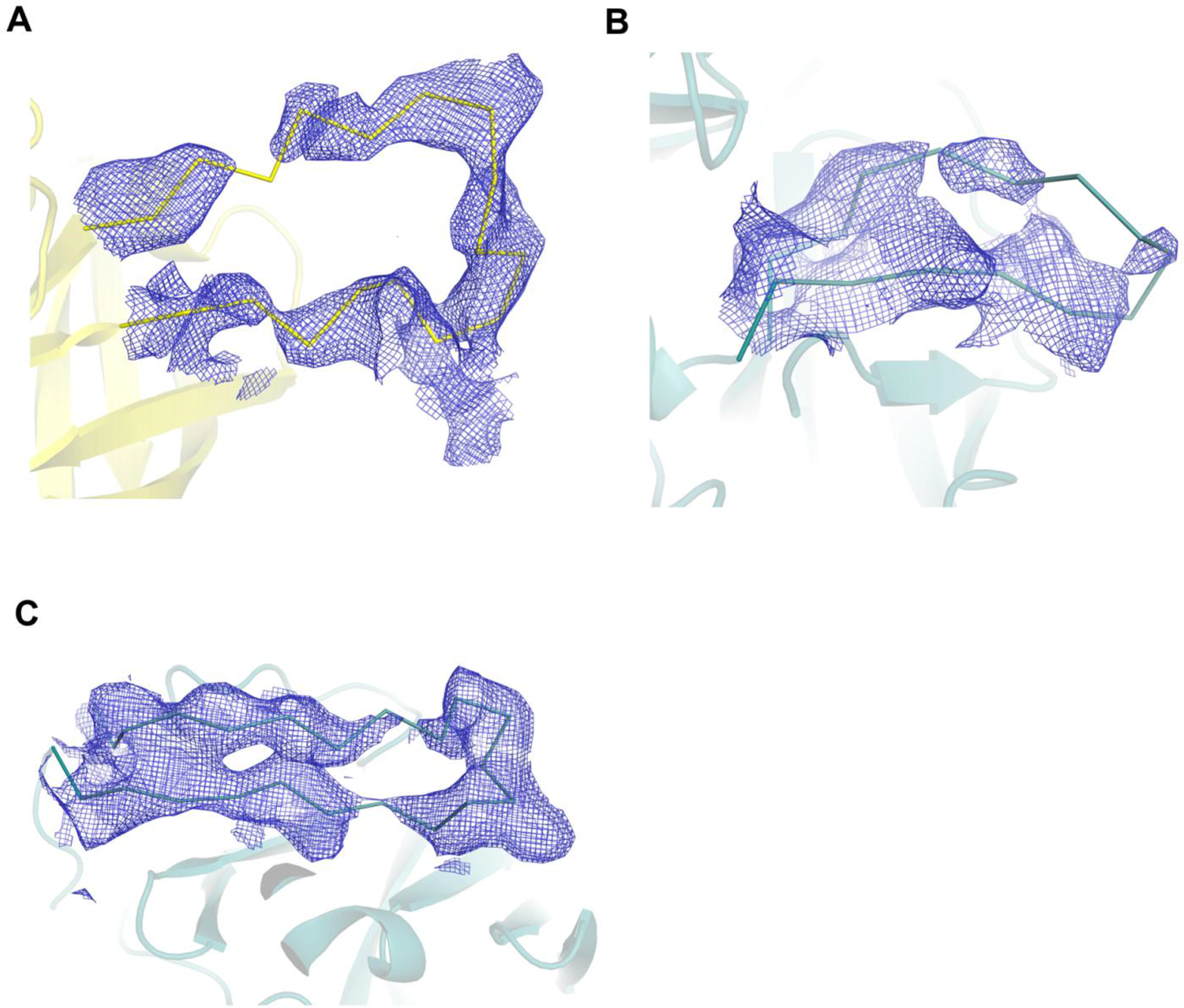
2Fo-Fc electron density maps of regions comprising different loops in ΔΔΔ. The maps for **(A)** LA loop (165-181 residues, contoured at 0.8 σ) from chain D, and the loops **(B)** L2 (320-333 residues, contoured at 0.8 σ) and **(C)** LD (255-275 residues, contoured at 1.0 σ) from chain A are modeled. LA loop is represented as yellow ribbon, while L2 and LD loops are represented as deep olive ribbons. The map is displayed as blue mesh.

